# Context specific ubiquitin modification of ribosomes regulates translation under oxidative stress

**DOI:** 10.1101/2024.05.02.592277

**Authors:** Shannon E. Dougherty, Géssica C. Barros, Matthew W. Foster, Guoshou Teo, Hyungwon Choi, Gustavo M. Silva

**Affiliations:** Department of Biology, Duke University, Durham, NC 27708, USA; Proteomics and Metabolomics Core Facility, Duke University, School of Medicine, Durham, North Carolina.NC 27701, USA; Department of Medicine, Yong Loo Lin School of Medicine, National University of Singapore, Singapore, Singapore; Singapore Lipidomics Incubator, Life Sciences Institute, National University of Singapore, Singapore

**Author notes:** Corresponding author: Gustavo M. Silva.

**Keywords:** ubiquitination, ribosome, translation control, stress response, proteomics, yeast

## Abstract

Cellular exposure to stress is known to activate several translational control pathways through ribosome ubiquitination. However, how unique patterns of ribosome ubiquitination act at the site-specific level to drive distinct modes of translation regulation remains unclear. To further understand the complexity of these ubiquitin signals, we developed a new targeted proteomics approach to quantify site-specific ubiquitin modification across the ribosome. This method increased the sensitivity and throughput of current approaches and allowed us to systematically measure the ubiquitin status of 78 ribosome peptides and ubiquitin linkages in response to stress. Using this method, we were able to detect the ubiquitination of several ribosome sites even in steady-state conditions, and to show that their modification increases non-stoichiometrically in a dynamic range of >4 orders of magnitude in response to hydrogen peroxide. Besides demonstrating new patterns of global ribosome ubiquitination, our study also revealed an unexpected increase of ubiquitination of ribosomal protein uS10/Rps20 and uS3/Rps3 independent of the canonical E3 ubiquitin ligase Hel2. Furthermore, we show that unique and mixed patterns of ribosome ubiquitination occur in a stress specific manner, depending on the nature of stressor and the enzymes involved. Finally, we showed that while deletion of *HEL2* further induces the integrated stress response in response to the nucleotide alkylating agent 4-NQO, deletion of the E2 conjugase *RAD6* leads to sustained translation only in response to H_2_O_2_. Our findings contribute to deciphering the complexity of the stress response at the translational level, revealing the induction of dynamic and selective ubiquitin codes, which shed light on the integration of important quality control pathways during cellular response to stress.

## INTRODUCTION

Cells have the remarkable ability to adapt to stress, which ensures their survival in ever-changing conditions. One way by which cells adapt is by implementing specific gene expression programs to combat stress (Gasch *et al*, 2000) (de Nadal *et al*, 2011). The induction of these gene expression programs has been extensively investigated at the transcriptional level (Gasch *et al*., 2000; Mascarenhas *et al*, 2008; Morano *et al*, 2012; Taymaz-Nikerel *et al*, 2016). However, discrepancies between mRNA and protein levels (Blevins *et al*, 2019; Buccitelli & Selbach, 2020; Vogel *et al*, 2011) emphasize translation as another influential regulatory mechanism in cellular stress. In response to various forms of stress, cells suppress global translation while simultaneously inducing the production of selective stress-response proteins (Barros *et al*, 2023; Costa-Mattioli & Walter, 2020; Gerashchenko *et al*, 2012; Shenton *et al*, 2006). This global translation suppression is crucial to favor the production of stress-related proteins while preventing damage to newly synthesized proteins, avoiding toxic gain-of-functions, and minimizing extraneous energy expenditure on unimportant proteins (Advani & Ivanov, 2019; Holcik & Sonenberg, 2005). Several molecular mechanisms, including the integrated stress response (ISR), are known to regulate translation initiation (Pakos-Zebrucka *et al*, 2016; Postnikoff *et al*, 2017) in cellular stress. However, how translation elongation is regulated is only beginning to be understood (Knight *et al*, 2020; Shenton *et al*., 2006). As the regulation of elongating ribosomes enables a rapid response to dynamic cellular conditions (Barros *et al*., 2023; Crawford & Pavitt, 2019), understanding the mechanisms underlying this regulation is vital to comprehend how cells respond and adapt to stressful conditions.

The redox-control of translation by ubiquitin pathway (RTU) is known to regulate translation elongation and support cellular stress adaptation in yeast (Dougherty *et al*, 2020; Silva *et al*, 2015). In response to oxidative stress, the RTU is activated and elicits a burst of ubiquitination causing elongating ribosomes to halt in a pre-translocation orientation (Zhou *et al*, 2020). In response to hydrogen peroxide (hereafter H_2_O_2_ or peroxide) stress, the deubiquitinating enzyme Ubp2 is inhibited (Santos *et al*, 2024), and the E2 ubiquitin conjugase Rad6 and E3 ligase Bre1 are involved in the modification of ribosomes with K63-linked polyubiquitin chains (K63-ub) (Silva *et al*., 2015; Simões *et al*, 2022). K63-ub is distinct from the more abundant K48-linked polyubiquitin (K48-ub) as it does not signal its target for proteasomal degradation (Manohar *et al*, 2019; Rahman & Wolberger, 2024; Xu *et al*, 2009). Rather, K63-ub has been shown to act as a regulatory signal in various biological processes, including DNA repair, intercellular trafficking, and immune signaling pathways (Erpapazoglou *et al*, 2014; Liu *et al*, 2018; Madiraju *et al*, 2022). Recent findings suggest that Rad6-mediated ubiquitination leads to ribosome pausing on isoleucine-proline motifs and contributes to mechanisms of translation repression (Meydan *et al*, 2023). However, several aspects of the underlying molecular mechanisms connecting ribosome ubiquitination and translational control in response to oxidative stress remain unclear.

In our previous work, we identified 78 lysine (K) residues distributed in 37 ribosomal proteins, referred to as ribosome ubiquitination sites, that showed over a 1.5-fold increase in Ub-modification under H_2_O_2_ compared to a yeast strain unable to build K63-ub chains (K63R mutant) (Back *et al*, 2019). However, it was unclear whether or to what extent these modifications accumulated in response to stress exposure. Further investigation into Ub-modified ribosomal sites has proven difficult due to the low abundance of ubiquitinated proteins relative to their non-ubiquitinated counterparts, methodological limitations of investigating each site independently, and the existence of multiple ubiquitin-mediated processes of ribosome regulation (Dougherty *et al*., 2020; Garshott *et al*, 2020; Inada, 2020). In one such process, the ribosome-associated quality control pathway (RQC), the E3 ligase Hel2 recognizes and ubiquitinates collided ribosomes (Matsuo *et al*, 2017; Matsuo & Inada, 2023), which recruits factors that dissociate the stalled ribosome from the mRNA transcript, leading to proteasomal degradation of the arrested peptide (Matsuo *et al*, 2023). Hel2 is known to directly modify or extend K63-ub chains of at least 5 sites in 4 different ribosomal proteins as part of the RQC and other decay pathways (Monem & Arribere, 2024). However, previous studies have shown that the increase in K63-ub observed in peroxide stress occurs independently of Hel2 (Silva *et al*., 2015; Simões *et al*., 2022; Zhou *et al*., 2020). As the RQC and the RTU can be induced under stress conditions (Silva *et al*., 2015; Yan *et al*, 2019), it is unclear how distinct ribosome ubiquitin signals coordinate the cellular response to stress and what their contributions are in varying cellular contexts.

In this study, we developed a new targeted proteomics method to quantify the dynamics of ribosome ubiquitination in response to oxidative stress in a relative and stoichiometric fashion. Using this approach, we identified 11 ribosomal sites that showed significantly increased ubiquitin modification in response to H_2_O_2_, including two previously known targets of Hel2. Using stable isotope-labeled internal standard peptides, we also showed that these sites are modified in a non-stoichiometric manner, ranging more than 4 orders of magnitude across different peptides. Moreover, we investigated the dependence of these ubiquitination events on distinct enzymes of the ubiquitin system, and the contribution of these pathways to global translation regulation. Our analysis of ribosome ubiquitination dynamics revealed that ubiquitination of unique ribosomal proteins such as uS3/Rps3 and uS10/Rps20 are differentially induced by distinct stressors, suggesting that enzymes such as Rad6 and Hel2 can compete for ribosome subpopulations. We also showed that the nature of oxidizers determines the antagonistic or synergistic impact on the ISR, resulting in additional layers of translation regulation. Our research highlights the importance of targeted proteomics methods to illuminate the quantitative complexity of the ubiquitin ribosomal code and further determine the role of this code in translation reprogramming under distinct environmental conditions.

## RESULTS

### Targeted proteomics uncovers the landscape of ubiquitin-modified ribosomes during H_2_O_2_

It has previously been shown that exposure to various stresses can lead to the ubiquitination of specific ribosomal sites (Back *et al*., 2019; Garshott *et al*., 2020; Matsuki *et al*, 2020; Yan *et al*., 2019). To investigate the role of these multiple sites in translation control, we first developed a Parallel Reaction Monitoring Mass Spectrometry (PRM-MS) based approach to detect and quantify abundances of ribosome Ub-modification with site-specificity (Fig. 1A). PRM is an ideal method to quantify post-translational modifications due to its selectivity, sensitivity, and mid throughput capabilities (Peterson *et al*, 2012). In PRM-MS, mass-to-charge (*m/z*) filtering reduces interference from non-target peptides, enhancing the capture and quantification of low-abundance species, such as those derived from ubiquitinated proteins (Ordureau *et al*, 2018; Peterson *et al*., 2012; van Bentum & Selbach, 2021). To perform PRM-MS, we designed 103 isotopically labeled (C-terminal Lysine (^13^C_6_,^15^N_2_) or Arginine (^13^C_6_,^15^N_4_)) tryptic reference peptides (JPT Peptide Technologies) (Table S1). These peptides corresponded to the previously identified ribosome sites (Back *et al*., 2019) and signature peptides representing each ubiquitin linkage type. Each synthetic peptide contains a di-Gly remnant (K-GG) at the site of the ubiquitinated lysine residue, consistent with the product of tryptic digestion of ubiquitinated proteins (Bustos *et al*, 2012; Kirkpatrick *et al*, 2005; Peng *et al*, 2003). Analysis of these heavy-labeled peptides serves as a guide to determine the peptide-specific retention times (RT) and *m/z* ratios that are needed to schedule data acquisition (van Bentum & Selbach, 2021). Moreover, the heavy-labeled peptides are spiked at 5 pM to serve as an internal standard for stoichiometric quantification of the biologically derived peptides (Kettenbach *et al*, 2011; Kito & Ito, 2008). Of the 103 synthetic peptides, 78 were successfully detected via data-dependent acquisition (DDA) MS and were used to generate the spectral library (MacLean *et al*, 2010) then used to schedule data acquisition and target the biologically derived peptides of interest (Table S1).

**Figure 1.**
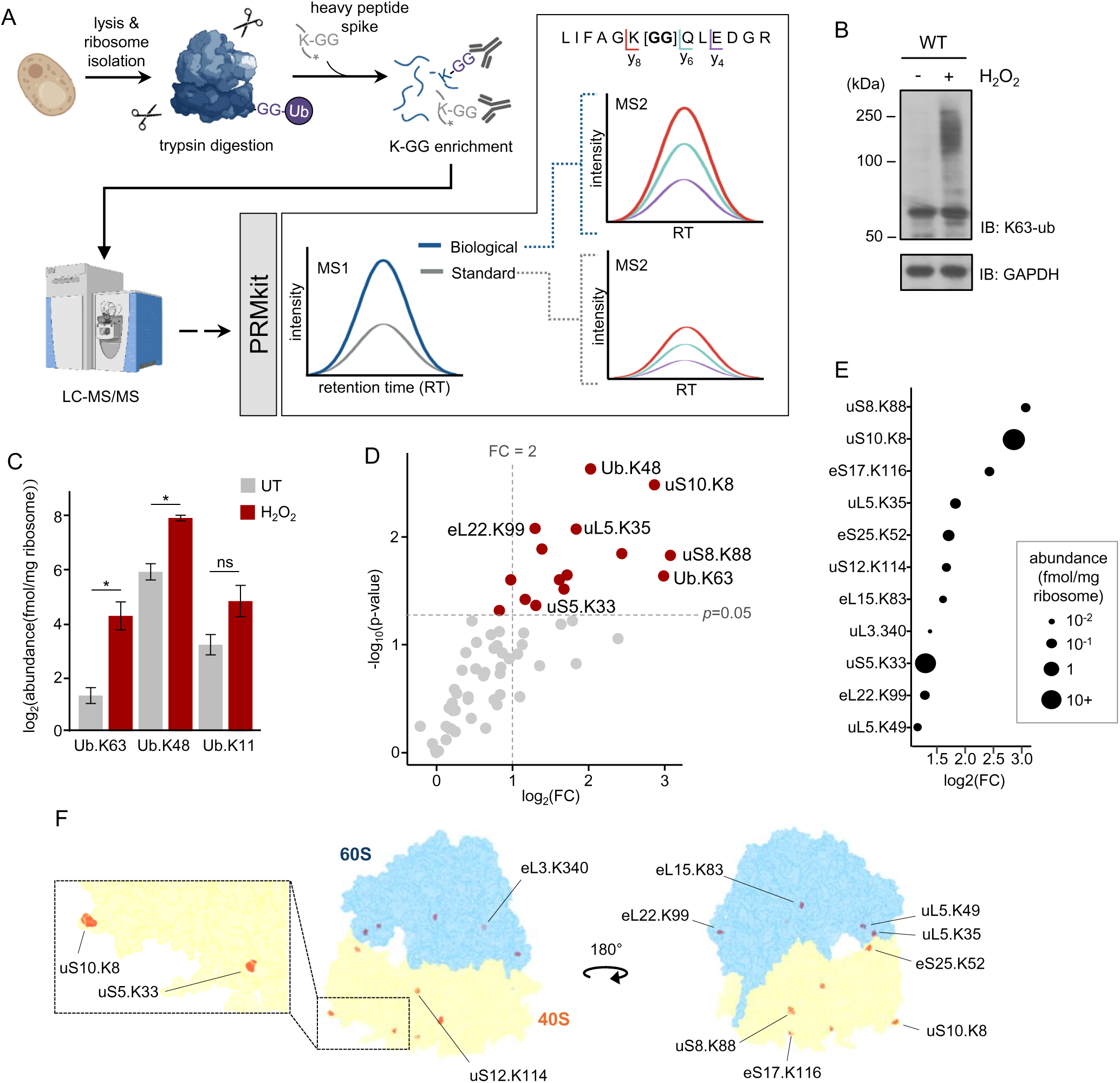
PRM-MS analysis of Ub-modified ribosome sites in H_2_O_2_. **(A)** Schematic of two-step protocol for ribosome isolation and enrichment of tryptic K-GG modified peptides. Heavy-labeled reference peptides were spiked in prior to enrichment with the anti-K-GG antibody. K-GG modified peptides were then analyzed by PRM-MS. PRMkit software was used to extract ion chromatograms and quantify peak area integration. **(B)** Western blot of K63-ub chains from cells treated in the presence or absence of 0.6 mM H_2_O_2_ for 30 minutes. anti-GAPDH was used as loading control. **(C)** PRM-MS quantification of signature ubiquitin linkage peptides. Plot shows abundance (fmol) of peptides per mg of ribosome from either untreated (UT) or 0.6 mM H_2_O_2_ treated (H_2_O_2_). Error bars represent SEM (n=5). Significance was calculated using a student’s paired *t* test, \**p* < 0.05 and *ns* is non-significant **(D)** Volcano plot of log_2_(fold change (FC)) of Ub-modified ribosome sites in untreated and H_2_O_2_ conditions. Significance was calculated using a student’s paired *t* test, \**p* < 0.05. **(E)** Plot of log2(FC) Ub-modified ribosome sites found to significantly increase from UT to H_2_O_2_ conditions (*p*<0.05) vs peptide abundance (fmol/mg of ribosome) calculated by normalization to heavy-labeled peptide. **(F)** Mapping of Ub–modified ribosome sites that showed statistically significant increase in H_2_O_2_ to the 3D structure of the 80S ribosome (PDB 6XIR) (Meng *et al*., 2023; Zhou *et al*., 2020). Inset shows the two most abundant sites K8 of uS10 and K33 of uS5 in the 40S subunit.

To quantify the site-specific ubiquitinated peptides that accumulate in response to peroxide stress, we applied PRM-MS to ribosomes from untreated and H_2_O_2_-treated cells. First, we harvested mid-log phase WT yeast cells before and after treatment with 0.6 mM H_2_O_2_. This H_2_O_2_ concentration is known to induce the accumulation of high levels of K63-ub chains (Fig. 1B). Following cell lysis, ribosomes were isolated by sucrose sedimentation and digested with trypsin to generate K-GG modified peptides (Fig. 1A). The heavy-labeled synthetic peptides were pooled and added to the digested samples. Using a K-GG remnant-specific antibody (PTMScan, Cell Signaling), we enriched for peptides derived from ubiquitinated proteins. These peptides were analyzed by PRM-MS using the pre-built spectral library from the heavy-labeled peptide analysis. To extract ion chromatograms and quantify peak area integration for all potential target-peptide transitions, we developed and implemented the PRMkit software (Fig. S1A-C). This tool offers a one-stop solution for PRM transition detection and peak integration guided by cross-sample consistency and reproducibility, thereby facilitating the selection of potential quantifier ions and downstream statistical analysis. Using PRMkit, we calculated the abundance of y-ions of targeted biological and heavy-labeled reference peptides across five pairs of untreated and H_2_O_2_-treated biological replicates. To determine the efficiency of this PRM-MS approach, we first evaluated its capacity to detect accumulation of ubiquitin linkages (K63-ub and K48-ub) known to be induced by H_2_O_2_ (Manohar *et al*., 2019; Silva *et al*., 2015). Analysis of the signature linkage peptide of K63-ub revealed an 8-fold increase upon treatment with H_2_O_2_ (Fig. 1C). Similarly, the signature linkage peptide for K48-ub increased around 4-fold in response to H_2_O_2_ treatment. Conversely, the K11-linked ubiquitin signature peptide did not increase significantly after H_2_O_2_ treatment, consistent with previous observations by immunoblot and MS-based approaches (Silva *et al*., 2015). These findings are consistent with previously published trends and highlight that our PRM-MS approach successfully captures H_2_O_2_ stress-induced changes in global ribosome ubiquitination.

Next, we set out to determine the changes in abundance of individual ubiquitinated sites in response to H_2_O_2_. Of the 78 detected peptides, 15, including K63-ub, K48-ub, and 13 ribosome sites, showed statistically significant increases in H_2_O_2_-treated compared to untreated conditions (Fig. 1D). Among these sites, 11 showed a greater than 2-fold increase due to H_2_O_2_ treatment (Fig. 1D). Noteworthy, all these sites were consistently detected in untreated cells, suggesting a basal level of ubiquitination of ribosomes even in steady-state, unstressed conditions. In addition to measuring the fold change of site-specific ubiquitination, our method allowed us to determine the overall abundance of each site modification through normalization to the heavy-labeled reference peptides. First, our data show that the abundance of these ribosome ubiquitinated peptides vary more than 4 orders of magnitude in a dynamic range of 2.2 amol to 72.5 fmol per mg of isolated ribosome (Table S2). When we consider only the 11 Ub-modified peptides with a greater than 2-fold increase under stress, they were quantified in a range of 0.004 to 72.5 fmol per mg of isolated ribosome after exposure to H_2_O_2_ (Fig. 1E, S2 A,B). Importantly, this increase in ubiquitinated peptides is likely driven by the accumulation of ubiquitin signals rather than increase of protein levels, as ribosomal proteins are known to have reduced expression during stress (An *et al*, 2020; Gerashchenko *et al*., 2012; Ghulam *et al*, 2020; Shenton *et al*., 2006). The two most abundant sites, uS10/Rps20 K8 and uS5/Rps2 K33, which were quantified to increase by about 7 and 2.5-fold in H_2_O_2_, respectively, are both located on the beak of the 40S ribosomal subunit, a region well-established as a hub of translational regulation (Fig. 1F) (Llácer *et al*, 2015; Meng *et al*, 2023; Wilson & Doudna Cate, 2012; Zhou *et al*., 2020). As specific Ub-modified sites undergo differential and non-stoichiometric modification, this finding suggests that a site-specific arrangement of ribosome ubiquitination, or Ub ‘code,’ arises in response to peroxide stress, which may have unique physiological implications.

Notably, uS10 K8 and uS5 K33 are reported as target sites for Ub-modification by Hel2 as part of the RQC pathway (Garshott *et al*., 2020; Matsuo *et al*., 2017; Yan *et al*., 2019). As Ub-modification of these sites increased under H_2_O_2_, our findings suggest that the RQC pathway could also be active in peroxide stress. However, the burst of K63-ub signals in H_2_O_2_ is not primarily mediated by Hel2 (Fig. 2A), (Silva *et al*., 2015; Simões *et al*., 2022), suggesting that Hel2 may only be modifying a particular subset of these ribosomal sites. To determine the role of Hel2 during H_2_O_2_ stress, we utilized MS to measure ribosome ubiquitination in *hel2Δ* cells (Table S3). The ubiquitin linkage signature peptides all exhibited similar trends in *hel2Δ* cells (Fig. 2B), as observed in wild type (WT), confirming that the major burst of ubiquitination in H_2_O_2_ occurs independent of Hel2. Although Ub-modification of most sites similarly increased under stress in *hel2Δ*, we observed no significant accumulation in Ub-modified uS5 K33 (Fig. 2C,D), further supporting that Hel2 mediates ubiquitination of this site (Garshott *et al*., 2020; Yan *et al*., 2019). However, we observed a partial Ub-modification of uS10 K8, which increases by ∼ 2-fold in H_2_O_2_-treated *hel2Δ*, about 30% of the increase observed in WT (Fig. 2C,D). This data indicates that, unlike uS5 K33, ubiquitination of uS10 K8 can be mediated by enzymes other than Hel2. Although not statistically significant, our method also detected increased ribosomal ubiquitinated sites in the absence of Hel2, such as eL30/Rpl30 K5. Ubiquitinated eL30 K5 increased ∼5x more upon stress in *hel2Δ* in comparison to WT (Fig. 2C). This finding suggests that reduction or suppression of ubiquitination of select sites (e.g. Hel2-dependent sites) can also activate compensatory responses, leading to new landscapes of ribosome ubiquitination. Our newly developed PRM-MS method allowed us to detect and quantify ubiquitin modification across more than 70 ribosome sites and revealed a complex code of site- and enzyme-specific Ub-modification induced upon H_2_O_2_ stress. However, it remains unclear what aspects of stress universally or selectively contribute to unique codes of site-specific ubiquitination of ribosomes.

**Figure 2.**
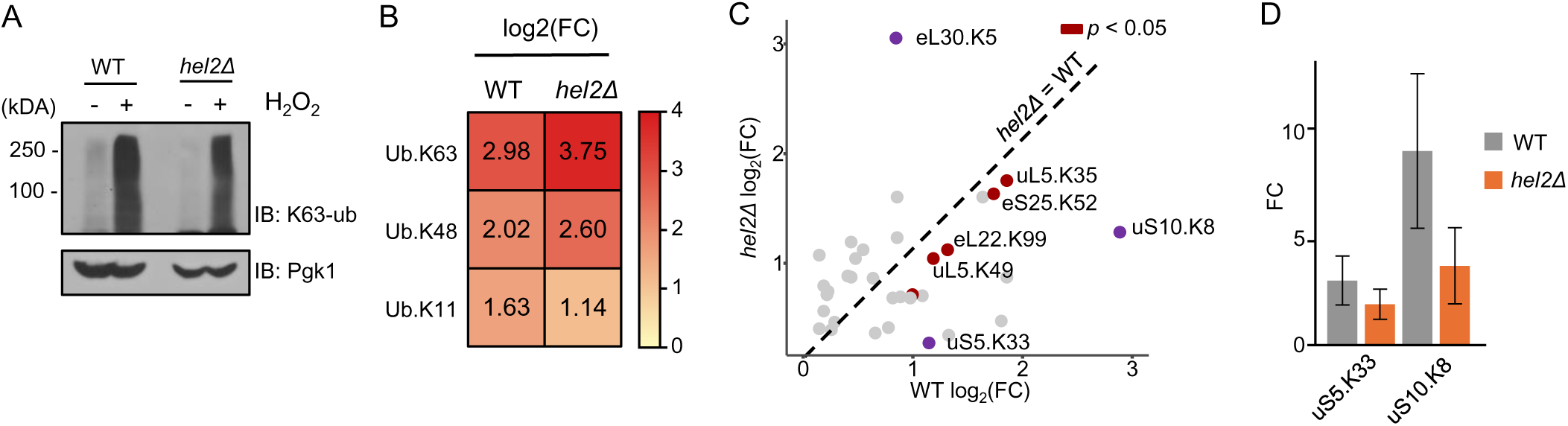
Quantification of ribosome site Ub-modification in the absence of Hel2. **(A)** Western blot of K63-ub chains from wild-type (WT) and *hel2Δ* cells treated in the presence or absence of 0.6 mM H_2_O_2_ for 30 minutes. anti-GAPDH was used as loading control. **(B)** Mass spectrometry comparison of fold change (FC) of main ubiquitin signature linkage peptides from untreated to H_2_O_2_ treated conditions in WT and *hel2Δ* strains. **(C)** Log_2_ FC plot of Ub-modified ribosome sites under H_2_O_2_ treatment for in WT (x-axis) and *hel2Δ* (y-axis). Significance was calculated using a student’s paired *t* test, \**p* < 0.05. **(D)** Comparison of FC of ubiquitinated uS10.K8 and uS5.K33 peptides due to H_2_O_2_ treatment in WT and *hel2Δ* strains.

### Pattern of ubiquitinated ribosomes is determined by the nature of the stressors

Our targeted proteomics analysis revealed a specific pattern of ribosomal ubiquitination that increases under peroxide stress. To validate our proteomics findings and further explore the mechanisms underlying the accumulation of Ub-modified sites during oxidative stress exposure, we investigated the ubiquitin status of select proteins. Using immunoblot, we examined the accumulation of ubiquitin-modified uS10 by expressing this protein endogenously tagged with the human influenza hemagglutinin (HA) (Matsuo *et al*., 2017). Indeed, we observed polyubiquitinated forms of uS10 upon treatment with low (0.6 mM) and high (5 mM) doses of H_2_O_2_ (Fig. 3A, lanes 2,3). Although we showed that uS10 can be ubiquitinated in response to H_2_O_2_ stress, polyubiquitinated uS10 has also been previously observed in response to the oxidizing agent 4-nitroquinoline 1-oxide (4-NQO) (Yan *et al*., 2019; Yan & Zaher, 2021). These findings suggest that ubiquitination of uS10 may be a universal feature of oxidative stress. To investigate this hypothesis, we measured the accumulation of K63-ub conjugates and the levels of polyubiquitinated uS10 upon treatment with low and high concentrations of 4-NQO (1 μg/mL, 5 μg/mL). These 4-NQO concentrations are known to induce stress that leads to lagged cell growth and ubiquitination of ribosomal proteins, respectively (Yan *et al*., 2019). We found that K63-ub accumulated heavily in 0.6 mM H_2_O_2_ but not in 5 mM H_2_O_2_ or either concentration of 4-NQO (Fig. 3A). However, immunoblot of uS10 in 4-NQO revealed polyubiquitination bands as observed in H_2_O_2_ stress (Fig. 3A, lanes 4,5), which increased with stress intensity. These findings indicate that while the global accumulation of K63-ub seems specific to peroxide stress (Back *et al*., 2019; Silva *et al*., 2015; Simões *et al*., 2022), ubiquitinated versions of particular proteins within this ribosomal code may accumulate across conditions. Previous studies into the mechanisms underlying the ubiquitination response in 4-NQO stress found the ribosomal protein uS3/Rps3 was ubiquitinated in conjunction with uS10 (Yan *et al*., 2019). The uS3 K212 ubiquitinated peptide was not detected via mass spectrometry, likely due to the size of the product peptide. Therefore, we used immunoblot to investigate whether the processes driving uS10 ubiquitination were also driving ubiquitination of uS3. Here, we observed mono-ubiquitinated uS3 (uS3-Ub_1_) upon exposure to 4-NQO but not H_2_O_2_ (Fig. 3B), indicating that uS3 might not be a main target of H_2_O_2_-induced ubiquitination. Importantly, following incubation of the cell lysate with the deubiquitinating enzyme USP2, we were able to observe the disappearance of the high molecular weight bands for uS10 and uS3, confirming that these bands, indeed, represent ubiquitinated versions of these proteins (Fig. S3A,B). Corroborating our proteomics data, these findings suggest that individual proteins may be ubiquitinated in a context-dependent manner and might serve unique roles in eukaryotic stress response.

**Figure 3.**
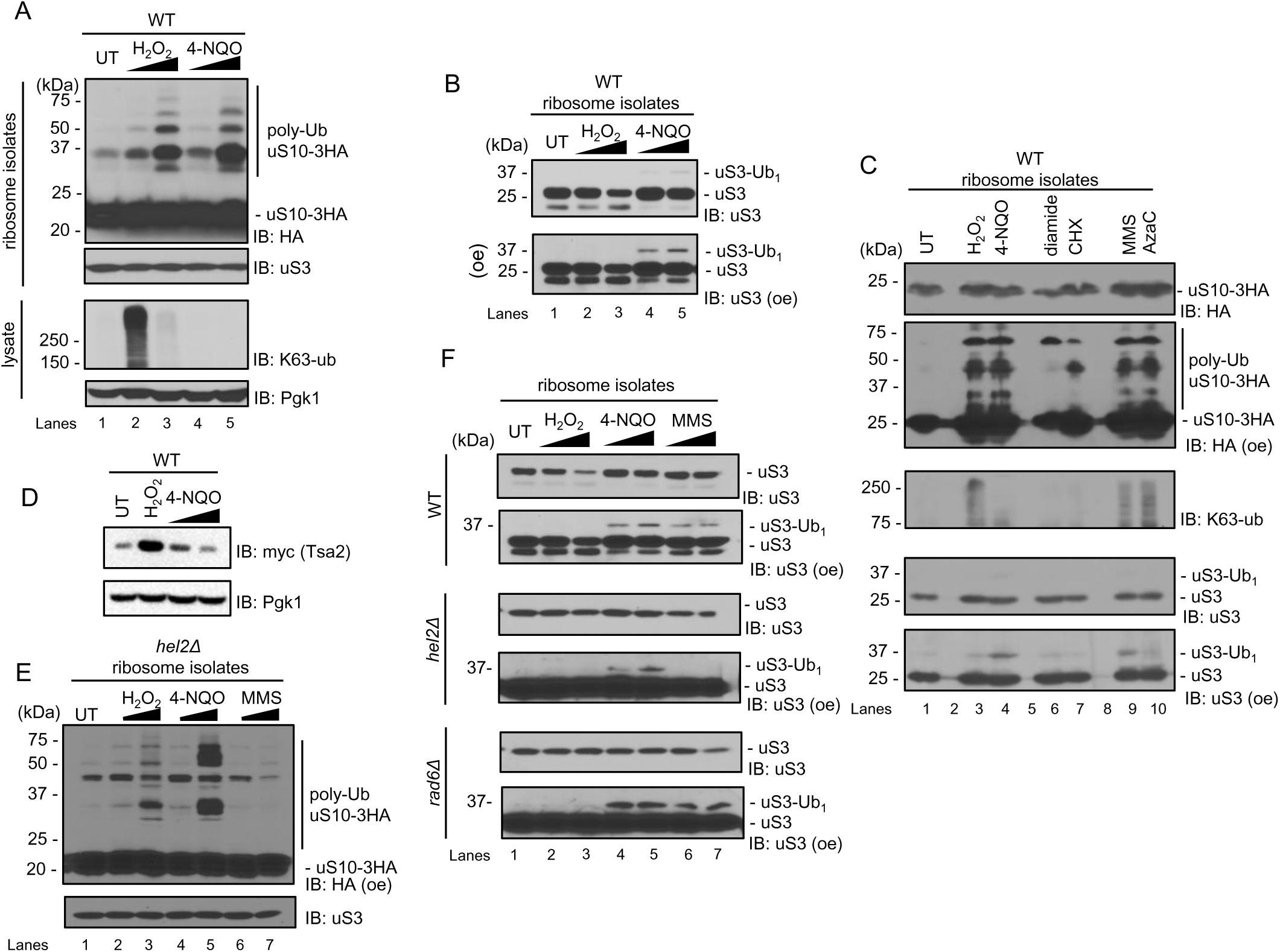
Dynamics of ribosomal protein ubiquitination in stress. **(A)** Western blot for K63-ub and uS10-3HA from lysate and ribosome isolate of WT cells in untreated (UT) and treated conditions. **(B)** Western blot of ribosomes isolated from UT and treated WT cells probed with anti-uS3. **(C)** Western blot for K63-ub, uS3, and uS10-3HA from ribosome isolate of WT cells in UT, 0.6 mM H_2_O_2_, 1 μg/mL 4-NQO, 1.5 mM diamide, 100 μg/mL cycloheximide (CHX), 0.1% methyl methanesulfonate (MMS), or 1 μM azacytidine (AzaC). **(D)** Western blot for expression of Myc-tagged Tsa2 in UT, H_2_O_2_ (0.6mM) and 4-NQO treated WT cells. **(E)** Western blot for polyubiquitinated forms of uS10 in ribosome isolates from *hel2Δ* cells in UT, and treated conditions. **(F)** Western blot for monoubiquitinated form of uS3 in ribosome isolates from WT, *hel2Δ,* and *rad6Δ* cells in UT and treated conditions. Unless otherwise noted, cells were treated with H_2_O_2_ (0.6 mM, 5 mM), 4-NQO (1 μg/mL, 5 μg/mL), and MMS (0.1%, 0.33%) concentrations for 30 min at 30°C. *oe,* overexposure. anti-Pgk1 was used as a loading control for lysate and anti-uS3 was used as loading control for ribosome isolation samples.

As both H_2_O_2_ and 4-NQO are considered oxidative stress agents, we set out to determine the features of stress that lead to uS10 and uS3 distinct ubiquitin patterns. To explore their unique ubiquitin landscapes, we used a series of oxidizing agents or chemical stressors known to induce ribosome ubiquitination (Silva *et al*., 2015; Simms *et al*, 2017; Yan *et al*., 2019). By using cycloheximide (CHX), a translation inhibitor that causes ribosome collisions and activates the RQC (Simms *et al*., 2017; Yan & Zaher, 2021), we show polyubiquitination of uS10 (Fig. 3C, lane 7), confirming that this ubiquitin accumulation is not an exclusive feature of oxidative damage. Ribosomal K63-ub chains did not accumulate globally under CHX (Fig. 3C, lane 7), reinforcing that peroxide-mediated reactions induce the formation of this linkage type. Further, we incubated yeast cells with diamide, a chemical that induces stress through the oxidation of thiol groups (Kosower & Kosower, 1995). However, diamide did not induce substantial amount of uS3-Ub_1_ and polyubiquitination of uS10 (Fig. 3C, lane 6). In agreement with the vast literature that discusses the unique biological and chemical properties of individual reactive oxygen species (ROS) and oxidizers (Jomova *et al*, 2023; Lushchak, 2014; Thannickal & Fanburg, 2000; Winterbourn, 2008), we show that the ribosomal Ub code identified in H_2_O_2_ stress is not a universal response to every oxidizing agent. Therefore, we hypothesized that oxidizing compounds such as H_2_O_2_ and 4-NQO might activate distinct ribosome ubiquitination patterns through their unique and main biochemical reactions. As H_2_O_2_ is a global oxidizer, we tested whether 4-NQO is also able to induce cellular oxidative stress. Here, we measured the levels of endogenously myc-tagged Tsa2 protein, an antioxidant protein whose expression is highly induced during oxidative stress by the redox regulation of a series of transcription factors (Wong *et al*, 2003; Wong *et al*, 2002). Interestingly, we found that Tsa2 was upregulated in response to H_2_O_2_, but not in response to 4-NQO (Fig. 3D). This indicates that despite similar levels of uS10 ubiquitination, 4-NQO does not induce cellular oxidative stress to the same extent as H_2_O_2_. These results suggest that the ribosomal ubiquitin code in 4-NQO is not due to a global oxidative stress function, as seen in H_2_O_2_, but rather through other oxidizing features of this chemical.

4-NQO is categorized as a chemical carcinogen due to its propensity to produce nucleotide adducts (Bailleul *et al*, 1989; Kohda *et al*, 1986; Yan *et al*., 2019). Therefore, we proposed that 4-NQO drives ribosome ubiquitination due to this selective oxidation of nucleotides. To test this hypothesis, we measured the ubiquitination of uS10 and uS3 in stress conditions known to cause nucleotide damage and disrupt RNA function, such as the alkylating agent methyl methanesulfonate (MMS) (Simms & Zaher, 2016; Yan *et al*., 2019). Upon exposure to MMS, we observed polyubiquitinated forms of uS10 and monoubiquitinated of uS3 (Fig. 3C), similarly to what was observed upon exposure to 4-NQO. Interestingly, exposure to DNA/RNA damage inducing agent 5-azacitidine (AzaC) (Aimiuwu *et al*, 2012; Bhuvanagiri *et al*, 2014; Khoddami & Cairns, 2013; Li *et al*, 1970), mostly produced polyubiquitination of uS10. These findings support a model in which the ribosomal Ub code is activated in a stress specific manner. As the ubiquitin pattern under RNA damage differs from the one observed in H_2_O_2_, these findings indicate that the ubiquitination system is highly sensitive and responsive to the nature of the stressor.

Since the ubiquitination system can respond to a variety of chemical stimuli, it poses the question whether the ubiquitination of ribosomes is carried systematically by the same enzymes. Previous studies have reported that Hel2 is responsible for the ubiquitination of uS10 and uS3 in 4-NQO and MMS stress (Nanjaraj Urs *et al*, 2024; Yan *et al*., 2019; Yan & Zaher, 2021). However, our proteomics data show partial ubiquitination of uS10 even in the absence of this E3 ligase (Fig. 2C,D). To validate our findings, we evaluated the pattern of uS10 ubiquitination in the *hel2Δ* strain. We confirmed that *HEL2* gene was deleted by PCR (Fig. S3C) and functional depletion of *HEL2* activity was further verified using an established translation stalling reporter GFP-6xCGA-FLAG-HIS3 (Matsuo *et al*., 2017). Using this reporter, we observed a strong ribosome arrest and rescue in the 6xCGA (poly-R) sequence in the WT and the expected bypass of the stalling sequence in the *hel2Δ* (Fig. S3D). After confirming *HEL2* deletion, we measured the ubiquitination of uS10 and uS3 in the *hel2Δ* background upon exposure to H_2_O_2_, 4-NQO, and MMS. Even in the absence of Hel2, we observed polyubiquitinated forms of uS10 in all conditions (Fig. 3E, S3E). This data corroborates our PRM-MS finding that the ubiquitination of uS10 is not exclusively mediated by Hel2 in H_2_O_2_ stress and suggests that this is also true in 4-NQO and MMS. Furthermore, we noted that polyubiquitinated uS10 increased with rising stress intensity for H_2_O_2_ and 4-NQO, but exhibited a different pattern with fewer high molecular bands under MMS. The difference between 4-NQO and MMS became more evident upon measurement of monoubiquitinated uS3 in cells lacking *HEL2*. Consistent with published findings, we observed a loss of monoubiquitinated uS3 (uS3-Ub_1_) in *hel2Δ* upon exposure to low (0.1%) and high (0.33%) concentrations of MMS (Fig. 3F, lanes 6, 7). In contrast, we only observed a partial loss of uS3-Ub_1_ in 4-NQO (Fig. 3F, lanes 4, 5), suggesting that, similar to uS10, the ubiquitination of uS3 is not exclusively mediated by Hel2 in a 4-NQO-specific context. Interestingly, we have previously shown that the majority of K63-ub signals that accumulate under H_2_O_2_ depends on the E2 Rad6 (Silva *et al*., 2015; Simões *et al*., 2022). Though, deletion of *RAD6* does not affect the pattern of uS3 ubiquitination under alkylating conditions (Fig. 3F). Altogether, these data indicate that the pattern of ribosome ubiquitination varies depending on the type of stress encountered and suggests that several enzymes of the ubiquitin system function in a complex manner to generate a stress-specific Ub-code.

### Rad6 and Hel2 distinctly impact broader translation stress response

Our analysis of ubiquitin dynamics revealed that the patterns of ribosome modification vary based on the nature of the stressors and the specific actions of E2 and E3 enzymes, such as Hel2 and Rad6. As translation is markedly inhibited under a variety of stress conditions (Williams & Rousseau, 2024), we set out to gain further insights into the impact of these enzymes in translation control. Here, we assessed translation through measurement of the incorporation of the methionine analog, homopropargylglycine (HPG) into newly synthesized proteins (Meydan *et al*., 2023). Here, we observed a 90% inhibition of translation in H_2_O_2_ and 75% inhibition in 4-NQO, indicating that exposure to both stressors suppresses translation (Fig. 4A). Next, we aimed to determine how Rad6 and Hel2 contribute to this global repression of translation. As we have previously observed (Meydan *et al*., 2023; Simões *et al*., 2022), deletion of *RAD6* significantly prevents translation repression that occurs in response to H_2_O_2_, however, we observed no significant difference in translation suppression in *hel2Δ* (Fig. 4B). This result suggests that the subset of ribosomal proteins modified by Hel2 is not critical to support the global repression of translation observed during H_2_O_2_. In turn, when we considered 4-NQO, depletion of Rad6 only had a minor effect on translation inhibition, but deletion of *HEL2* led to a significant further 20% reduction in protein synthesis (Fig. 4B). In agreement with what has been proposed before (Yan *et al*., 2019; Yan & Zaher, 2021), Hel2 seems to act by alleviating the translation repression specifically in the context of 4-NQO stress.

**Figure 4.**
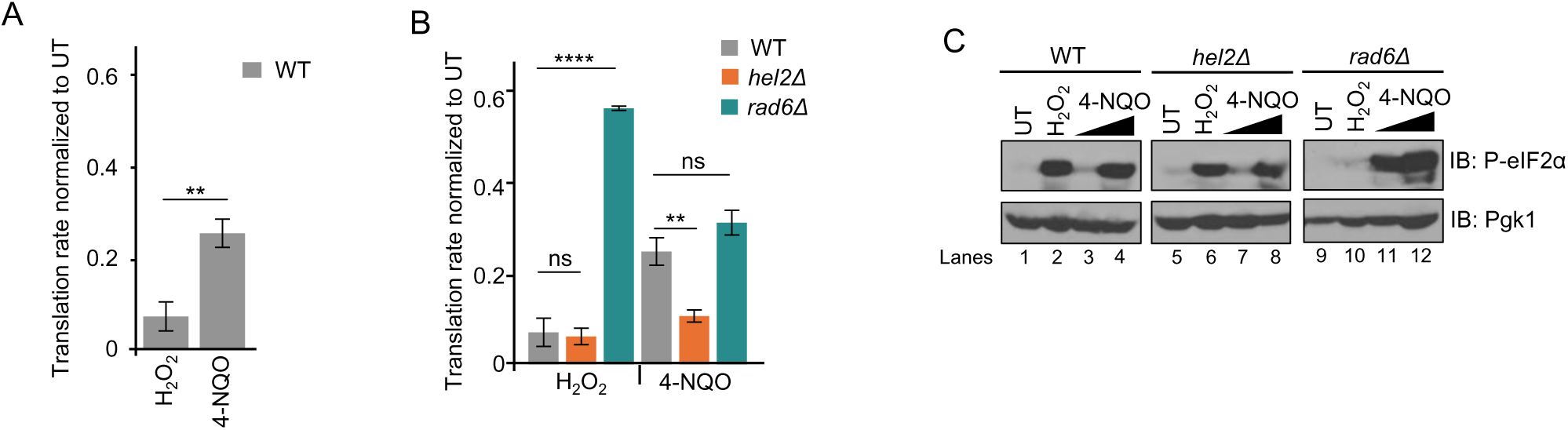
Stressor-specific suppression of global translation. **(A-B)** Quantification of HPG incorporation upon exposure to 0.6 mM H_2_O_2_ and 1 μg/mL 4-NQO in wild-type (WT), *hel2Δ*, and *rad6Δ* strains normalized to untreated (UT). Error bars represent SEM (n=2). Significance was calculated using Tukey’s multiple comparison test, \*\**p*< 0.005, **** *p<* 0.0001. *ns,* non significant **(C)** Western blot of phosphorylated eIF2α (P-eIF2α) from WT, *hel2Δ*, and *rad6Δ* cells in UT, 0.6 mM H_2_O_2_ and 4-NQO (1 μg/mL, 5 μg/mL). Anti-Pgk1 used as a loading control

The suppression of translation in response to cellular stress is often attributed to the activity of the Integrated Stress Response pathway (ISR) in reducing the initiation step of translation (Pakos-Zebrucka *et al*., 2016). In the yeast ISR, translation initiation factor eIF2α is phosphorylated by the Gcn2 kinase (Postnikoff *et al*., 2017). The phosphorylated eIF2α (P-eIF2α) then inhibits translation initiation by preventing the formation of active ternary complex (eIF2-GTP-tRNA_i_^Met^), which results in suppression of global translation (Mascarenhas *et al*., 2008; Pakos-Zebrucka *et al*., 2016). Therefore, we asked whether the enzyme-dependent effects on translation observed in *hel2Δ* and *rad6Δ* strains may be due to differential induction of the ISR pathway. By employing an antibody specific to the phosphorylated version of eIF2α (P-eIF2α), we observed in the WT a substantial increase of P-eIF2α upon treatments with H_2_O_2_ and the high 4-NQO concentration (Fig. 4C, lanes 2,4). Similar trends were observed in *hel2Δ*, however, a contrasting pattern was observed in the absence of Rad6. In *rad6Δ,* we observed low levels of P-eIF2α upon treatment with H_2_O_2_, which could explain the lack of translation repression observed (Fig. 4B). However, we observed a considerable increase in P-eIF2α upon treatment with 1 μg/mL 4-NQO (Fig. 4C, lane 11), which does not reflect in increased repression of global protein synthesis. Though, these findings further suggest that Rad6 and Hel2 also interacts with the ISR in inverse modes and in a stress-specific fashion.

To further understand the disconnect between high P-eIF2α levels and active translation, we investigated whether the ISR is being fully activated by measuring the gene products regulated by this pathway. Phosphorylation of eIF2α by Gcn2 reduces translation re-initiation, which allows for the synthesis of the transcription factor Gcn4 and the expression of its regulon (Hinnebusch, 2005; Pakos-Zebrucka *et al*., 2016). Thus, to understand Gcn4 activation, we investigated RNA-seq datasets for *hel2Δ* exposed to 0.1% MMS and *rad6Δ* exposed to 0.6 mM H_2_O_2_ (Meydan *et al*., 2023; Yan & Zaher, 2021). As MMS promotes ribosome ubiquitination through analogous mechanisms as 4-NQO, we used this condition as a proxy for RNA stress. As previously observed (Yan & Zaher, 2021), 124 genes of the Gcn4 regulon are induced >1.5 fold in the WT under MMS and even further in the (167, <1.5 fold) in *hel2Δ* cells (Fig. 5A,D). However, the Gcn4 regulon is only mildly activated in the presence of 0.6 mM H_2_O_2_. This analysis revealed that only 7 genes of the Gcn4-regulon showed increased transcript levels (> 1.5) in H_2_O_2_ stress in the WT compared to MMS (Fig. 5B). Of these 7 genes, only the catalase *CTT1*, an antioxidant enzyme that decomposes hydrogen peroxide (Martins & English, 2014), seems to be exclusively expressed under H_2_O_2_. The number of genes of the Gcn4-regulon upregulated jumps to 65 in *rad6Δ* (Fig. 5C), but it is still inferior to the 167 genes upregulated in *hel2*Δ under MMS treatment. Therefore, this analysis displays that not only translation and ubiquitination respond to stress in unique ways, but distinct Gcn4-mediated responses are also observed at the gene expression level as part of the ISR (Fig. 5D). To systematically compare ISR activation across conditions, we used a *GCN4-*LacZ reporter (Dever, 1997; Meydan *et al*., 2023), which contains the *GCN4* coding sequence with native uORFs (Hinnebusch, 2005) and LacZ becomes expressed through the activation of the ISR. We showed that ISR levels are enhanced in both strains in untreated conditions, suggesting that deletion of *RAD6 and HEL2* enhance the basal levels of cellular stress (Fig. 5E). As expected, Gcn4-LacZ levels are increased in *hel2Δ* treated with 4-NQO but not under H_2_O_2_. Surprisingly, the activation of ISR did not change in *rad6Δ* upon treatment with 4-NQO (Fig. 5E, S4A) regardless the high levels of P-eIF2α (Fig. 4C). Thus, our findings further highlight a complex relationship of the ISR with Hel2 and Rad6, where Hel2 dampens the activity of the ISR, but Rad6 is required to support translation repression under H_2_O_2_. This also suggests that the regulation promoted by Rad6 plays a dominant role and translation is not fully inhibited in *rad6Δ* despite the levels of P-eIF2α.

**Figure 5.**
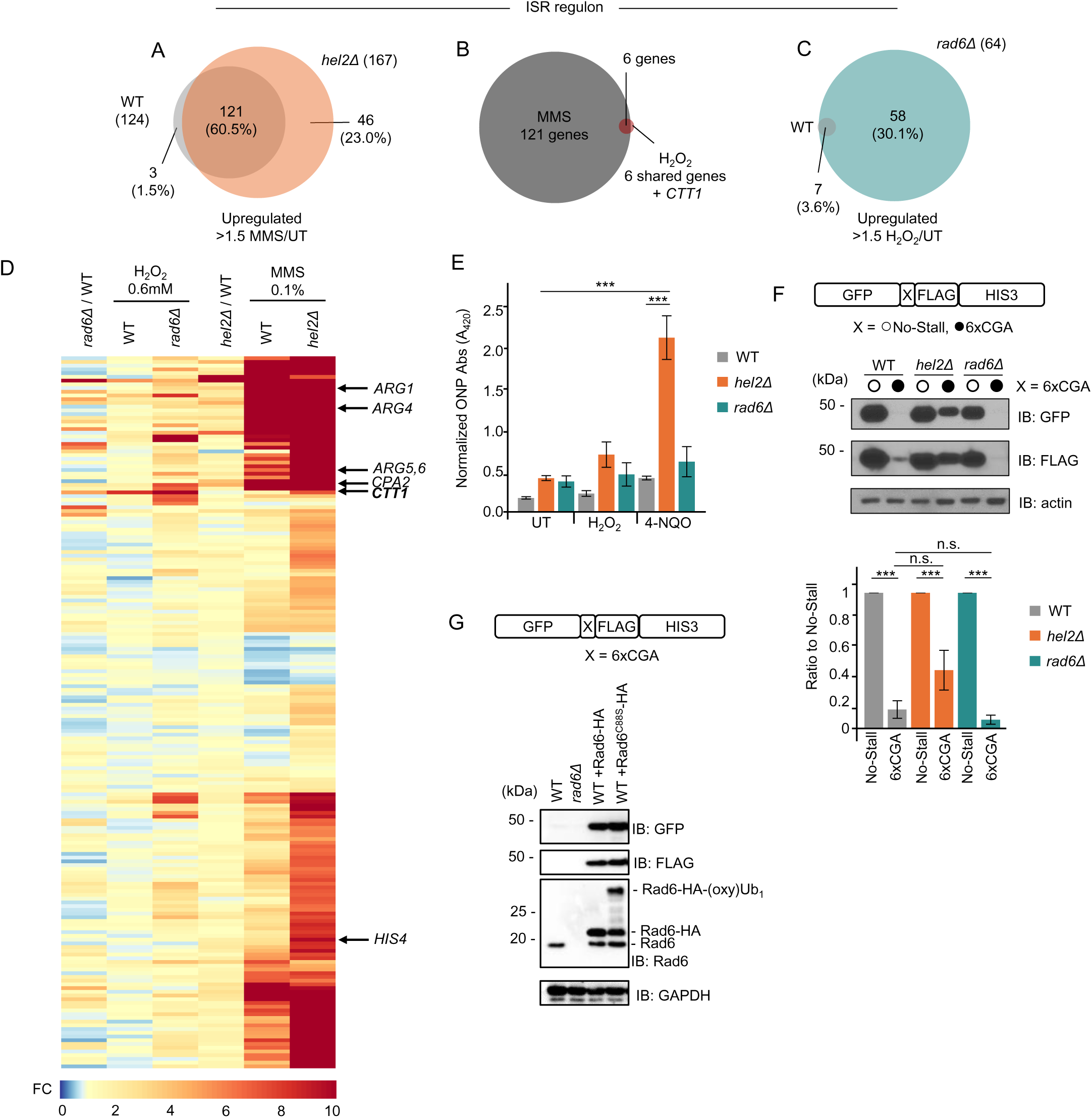
Enzyme- and stress-specific induction of *GCN4* pathway. **(A-C)** Venn diagram of RNA-seq data showing distribution of *GCN4* regulon that are upregulated (fold change >1.5) upon treatment with (**a**) 0.1% MMS in WT and *hel2Δ,* and (**b**) shared with WT treated with 0.6 mM H_2_O_2_. (**c**) Genes upregulated under 0.6 mM H_2_O_2_ in WT and *rad6Δ*. **(D)** Heatmap of RNA-seq meta-analysis of known GCN4-targeted stress response transcripts. RNA-seq data from GSE150790 (Yan & Zaher, 2021) and Data from metanalysis of RNA-seq analysis from GSE150790 (Yan & Zaher, 2021) and GSE226082 (Meydan et al., 2023). **(E)** Reporter assay for Gcn4 activity by *GCN4-lacZ* fusion construct in WT, *hel2Δ,* and *rad6Δ*. Cells were treated for 2 hours with 0.6 mM H_2_O_2_ or 1 μg/mL 4-NQO before protein extraction. ONPG absorption values detected at 420 nm were normalized to protein levels. Error bars represent SEM (n=2). Significance was calculated using Tukey’s multiple comparison test, *** *p*< 0.001. **(F)** Western blot anti-GFP and anti-FLAG for full-length GFP-X-FLAG-HIS3 reporter expressing No-Stall (open circle) and 6xCGA (full circle) sequences in WT, *hel2Δ*, and *rad6Δ*. Anti-actin was used as a loading control. Anti-FLAG was quantified and 6xCGA levels were normalized to strain-specific expression of the No-Stall reporter. Error bars represent SEM (n=3). Significance was calculated using Tukey’s multiple comparison test, \*\*\**p* < 0.001. **(G)** Western blot anti-GFP and anti-FLAG for full-length GFP-6xCGA-FLAG-HIS3 reporter in WT, *rad6Δ*, and WT strains expressing HA-tagged Rad6, or Rad6^C88S^. Levels of Rad6 measured by anti-Rad6 and anti-GAPDH was used as a loading control.

Finally, to further understand the relationship between Rad6 and Hel2, we investigated whether Rad6 impact Hel2 function in the RQC pathway. Using the GFP-6xCGA-Flag-HIS3 reporter, we observed that the absence of Hel2 leads to bypassing of the stalling sequence and partial production of the full-length product (Fig. 5F). Deletion of *RAD6* led to an enhanced stalling compared to the WT cells, suggesting the absence of Rad6 allowed for a more effective RQC function. We thus hypothesized that Hel2 and Rad6 could be competing for ribosome surfaces as part of their surveillance mechanisms. Thus, we episomally overexpressed Rad6 and determined the levels of full-length product from the stalling reporter. Corroborating our hypothesis, expression of WT Rad6 led to increased reporter production (Fig. 5G, S4B). Interestingly, expression of the catalytic dead Rad6 C88S also led to increased bypassing of the stalling sequence, suggesting that Rad6 ubiquitination activity is not required to compete with Hel2. Thus, our findings contribute to a model in which an intrinsic stoichiometric balance between Rad6 and Hel2 contributes to the outcomes in translation and quality controls pathways in response to specific stress conditions.

## DISCUSSION

Ribosome ubiquitination is a critical feature of translation reprogramming, particularly in conditions of cellular stress. Ubiquitination has been implicated in the rescue of stalled ribosomes (Matsuo *et al*., 2017), translation pausing (Zhou *et al*., 2020), in addition to processes of ribosome synthesis and degradation (Dougherty *et al*., 2020; Martín-Villanueva *et al*, 2021; Martínez-Férriz *et al*, 2022). This diversity of signals is determined by a complex code, which involves individual ribosome sites, enzymes of the ubiquitin system, polyubiquitin chain linkages, and specific cellular conditions. However, how these signals change in dynamic environments and how these Ub-mediated events are integrated into the intricate fabric of translation reprogramming under stress is only beginning to be elucidated. Here, we developed a targeted proteomics method (Fig. 1A) to simultaneously detect and quantify changes in ubiquitination across numerous known ribosomal sites. Using this method, we defined a broad landscape of ribosome ubiquitination in response to H_2_O_2_ and demonstrated how dynamic this process can be (Fig. 1D, 1E). The sensitivity and quantitative nature of our method allowed us to observe basal levels of ubiquitination that are usually undetected by standard antibody-based methods. This suggests that regulatory ubiquitination of ribosomes is more dynamic than previously thought, and functional cycles of ubiquitination and deubiquitnation are likely prevalent in unstressed eukaryotic cells. Stress might disrupt these cycles by regulating selective enzymes (Santos *et al*., 2024; Sheng *et al*, 2024; Simões *et al*., 2022) or by increasing the levels of available substrates (Chen *et al*, 2023; Iyer *et al*, 2023; Vind *et al*, 2020), such as stalled ribosomes, leading to the accumulation of ubiquitinated targets. The non-stoichiometric accumulation of these ribosome ubiquitinated sites (Fig. 1D, 1E, S2) also indicates that pathways are activated at different levels affecting unique numbers of ribosome particles. A combination of proteomics and immunoblot also revealed that Ub-modified ribosome sites known to be substrates of the E3 Hel2, can also be modified by other enzymes. The pattern of ubiquitinated sites is defined in a stress-specific manner, and our results continue to move us away from a model where one stress condition only activates a single pathway, regulated by a single enzyme, and well-defined set of substrates (Fig. 2C, 3C, 3F). Analogous to the histone code, where histone N-terminal tails can be modified by several PTMs that define the activation or silencing of transcription (Bowman & Poirier, 2015; Cavalieri, 2021), we propose that combinations of ubiquitin marks define the nature and magnitude of downstream pathways that are activated in response to individual stresses.

In response to H_2_O_2_, ubiquitination leads to ribosome pausing at the pre-translocation stage of elongation (Zhou *et al*., 2020) with a preference for isoleucine-proline encoding mRNA motifs (Meydan *et al*., 2023). However, the mechanisms driving this pause is unclear. Our structural work has suggested that K63-ub chains that accumulate upon H_2_O_2_ destabilize the ribosomal P-stalk, which could arrest elongating ribosomes by preventing the interaction or GTPase activity of the translation elongating factor eEF2 (Zhou *et al*., 2020). As we observed a non-stoichiometric change of ubiquitin sites, a model that all ribosomal sites must be modified to affect eEF2 function seems unlikely. Characterization of the landscape of ribosome ubiquitinated sites that are modified in response to stress independently of Hel2 provides new avenues for research into how this site-specific ubiquitination mediates RTU-related ribosome function. Notably, several of these sites are localized to the tRNA exit tunnel and thus serve as ideal candidates for further investigation into the mechanisms underlying translational control. Also of interest, we detected ubiquitination at known RQC sites in absence of Hel2 (Fig. 2C, 3E, 3F), introducing the possibility that a partially compensatory system exists for the RQC or that the single ribosome site can be a target for modification by multiple pathways based on the biological context in which it occurs.

In our proteomics analysis, we identified two H_2_O_2_-induced Ub-modified ribosome sites known to be dependent on Hel2 (Fig. 2C) and induced by 4-NQO oxidative stress (Yan *et al*., 2019). Through analysis of ubiquitination dynamics, we were able to determine that ubiquitination of uS10 and uS3 during 4-NQO and H_2_O_2_ stress are induced by different mechanisms. Here we highlight that different chemicals might change the pattern, quantity, or even the length of accumulated ubiquitin signals (Fig. 3A, 3C, 3F). These findings illustrate the granularity and responsiveness of the ubiquitination system to stresses, even when they would be considered variations of the same class of stress, such as oxidative stress. Our data show that for each unique stress condition, a complex model of sensors, writers, readers, and erasers are at full display on translation control. Therefore, further characterization of these molecular players must be done to fully dissect the complex mechanisms of stress-induced ubiquitin-mediated translational control. It is not farfetched to speculate that a fraction of ribosomes that are stalled in response to oxidative stress could lead to collisions and activation of RQC. Supporting this notion, our Disome-seq analysis under H_2_O_2_ (Meydan *et al*., 2023) revealed a similar pattern of isoleucine-proline pausing as observed for arrested monosomes. However, the sheer number of collisions would likely be inferior to what could be observed under mild translation inhibition such as in the presence of cycloheximide or anisomycin (Wu *et al*, 2020; Yan & Zaher, 2021). Interestingly, we also show that these pathways and enzymes likely interact and compete for ribosome substrates (Fig. 5F, 5G), which adds to the fluidity of these pathways. Therefore, improved quantitative methods must continue to be implemented to dissect the complexity and outcomes of these ubiquitin codes.

Our investigation into the contributions of the E2 Rad6 and the E3 Hel2 to the cellular response to stress revealed two distinct relationships with the ISR: Rad6 is critical for translation repression in response to H_2_O_2_, while Hel2 alleviates the impact of the ISR in translation inhibition during 4-NQO stress (Fig. 4B, 5E). The effect of this Hel2 mitigation of ISR was previously described (Nanjaraj Urs *et al*., 2024; Yan & Zaher, 2021) and here we show that it is stress specific. Interestingly, the impact of Rad6 or Hel2 in the ISR was not observed in the higher concentration of 4-NQO stress (Fig. S4A), indicating that in addition to the nature of the stressor, its intensity also defines which pathways are functional and predominant. In this sense, the RQC and the RTU might act upstream of the ISR at the elongation level under moderate and specific stress conditions. These findings suggest that RTU induction in peroxide stress acts as an early defense mechanism to shut down translation before prolonged arrest or damage can occur. Alternatively, induction of the RQC functions is an attempt to mitigate the effects of already existing arrests, reducing the impact of the ISR in repressing initiation. In the absence of Hel2, the ISR becomes the main active pathway, leading to enhanced inhibition of initiation. Upon acute stress conditions, mechanisms of elongation control are not needed or functional as initiation is largely repressed and thus, deletion of *RAD6* or *HEL2* do not affect the overall levels of translation repression (Fig. S4A). Therefore, understanding the expression and activity of these enzymatic systems and the status of translation will allow us to develop integrated models that take into account several mechanisms of control that occur simultaneously in individual subpopulations of ribosomes.

### Data availability

The LC-MS/MS proteomics data (.RAW files) have been deposited to the PRIDE repository partnered with ProteomeXchange with the dataset identifier PXD056496.

PRMkit software can be found here: https://github.com/PRMkit/PRMkit

## Supporting information

This article contains supporting information.

## Acknowledgments

We thank the Duke Proteomics and Metabolomics Shared Core Facility for support with mass spectrometry data acquisition. We are indebted to Toshifumi Inada for kind donation of plasmids and yeast strains.

## Author contributions

G.M.S. conceived, supervised, and funded the study. S.D., G.C.B., and M.F., generated resources and performed experiments. S.D., G.C.B., M.F., H.C., T.G., and G.M.S. analyzed data. S.D. and G.M.S. wrote the manuscript. All authors edited and contributed to its final form.

## Funding and additional information

This work was supported with funds from NIGMS R35 Award and the Chan Zuckerberg Initiative (GM137954 and SDL2022-253663, respectively, to G.M.S.). The content is solely the responsibility of the authors and does not necessarily represent the official views of the National Institutes of Health. The work was also supported in part by Singapore Ministry of Education (T2EP20223-0010), National Research Foundation and Agency for Science, Technology and Research (I1901E0040) and National Medical Research Council (CG21APR1008). Acquisition of the ThermoFisher Fusion Lumos mass spectrometer was supported in part by the Office of The Director, National Institutes of Health of the National Institutes of Health under Award Number S10OD024999.

## Conflict of interest

The authors declare that they have no conflicts of interest with the contents of this article.

## MATERIALS AND METHODS

### Yeast strains, plasmids, and growth conditions

#### Microbe strains

All *Saccharomyces cerevisiae* strains used in this study are listed in Table S4. Unless otherwise noted, yeast strains were grown in synthetic defined (SD) medium composed of D-Glucose (BD Difco, #215510), yeast nitrogen base (BD Difco, #291940), and required amino acids at 30°C and 200 rpm agitation. This study used the *E. coli* strain NEB10-beta (New England Biolabs #C3019H) grown in LB-medium (Sigma-Aldrich #L3022) at 37°C and 200 rpm agitation.

#### Strain generation

Standard recombination methods were used to delete and tag genes and were confirmed by PCR. Plasmids were constructed using either T4 ligation (New England Biolabs #M0202L) or HiFi DNA Assembly methods (New England Biolabs, #E2621S). All plasmids used in this study are listed in Table S5.

#### Growth conditions

For all mass spectrometry experiments, SUB280 and SUB280 *hel2Δ* yeast were grown in synthetic defined (SD) medium (BD Difco, #215510, BD Difco, #291940) and drop-out amino acid medium without Leu and Trp (Sigma, #Y0750). Follow up experiments in SUB280 and W303 derivatives were grown in SD complete media by using dropout amino acid supplements without Ura (Sigma, #Y0751) and supplementing back with Uracil (Sigma, #U0750). SUB62 strain derivatives were grown in (SD) medium composed of D-Glucose (BD Difco, #215510), yeast nitrogen base (BD Difco, #291940) and drop-out amino acid medium without Ura (Sigma, #U0750). Unless specified, yeast cells were cultivated at 30°C at 200 rpm agitation to mid-log phase. Unless otherwise stated, yeast cells were treated for 30 minutes with the following stressors/concentrations: freshly diluted H_2_O_2_ (peroxide) (Sigma, #216763) was added to a concentration of 0.6 mM or 5 mM, 4-nitroquinoline 1-oxide (4-NQO) (Sigma-Aldrich #N8141) was added to a concentration of 1 μg/mL or 5 μg/mL, Methyl methanesulfonate (MMS) (Sigma-Aldrich #129925) was added to a concentration of 0.1% or 0.33% v/v.

### Yeast protein extraction and western blots

#### Yeast protein extraction

Cells were disrupted by glass-bead agitation at 4°C in standard lysis buffer containing 50 mM Tris-acetate pH 7, 100 mM NaCl, 30 mM MgCl2, 20 mM iodoacetamide (IAM), and 1X protease inhibitor cocktail set I (Sigma, #539131). Cells intended for sucrose sedimentation were disrupted in standard lysis buffer with 1X protease inhibitor cocktail replaced by 1x complete mini EDTA-free protease inhibitor cocktail (Roche #04693159001). 1X Phosphatase Inhibitor (Sigma-Aldrich #P5726) was added to the standard lysis buffer for samples probed for phosphorylated proteins by western blot. Extracts were clarified by centrifugation, and protein concentration was determined via Bradford Assay (Bio-Rad, #5000205).

#### Ribosome sucrose sedimentation

Yeast lysates were sedimented by ultracentrifugation for 120 min at 70,000 rpm (Beckman Optima Max-TL, TLA-110 rotor) at 4°C in a 50% sucrose cushion buffered in 50 mM Tris-acetate, pH 7.0, 150 mM NaCl, and 15 mM MgCl2. Ribosome pellet was resuspended in the same lysis buffer and protein concentration was determined via Bradford Assay (Bio-Rad, #5000205) prior to western blot.

#### Western blot analysis

Proteins were separated by standard 10-15% SDS-PAGE and transferred to PVDF membrane (ThermoFisher, #88518). Immunoblotting was performed using the antibodies in Table S6. Western blots were quantified using ImageJ Software (Schneider *et al*, 2012), and Student’s t-test was used to calculate statistically significant differences with a p-value cutoff of 0.05.

### Mass spectrometry experiments

#### Heavy labeled reference peptide synthesis and characterization

Heavy labeled reference peptides (Table S1) for PRM containing isotopically labeled C-terminal Lysine (13C6,15N2) or Arginine(13C6,15N4) residues were synthesized by JPT Peptide Technologies. Heavy labeled reference peptides were reconstituted in 100 mM ammonium bicarbonate (AmBic) to a final concentration of 100 pmol/uL. To generate a spectral library, 200 fmol of stable isotope-labeled peptides were analyzed by LC-MS/MS using a Waters M-Class interfaced to a Thermo Fusion Lumos MS/MS. Liquid chromatography utilized a trap-elute configuration and 90 min gradient. MS/MS used data-dependent acquisition with an orbitrap MS2 resolution of 30,000, AGC target of 5E4, max injection time (IT) of 54 ms and stepped normalized collision energy (NCE) of 30 -/+ 5%. Mass spectra were imported to Skyline (MacLean *et al*., 2010) to build a spectral library using the MS-Amanda search engine.

#### Yeast ribosomal peptide preparation

SUB280 WT or SUB280 *hel2Δ* yeast were grown to mid-log phase in SD medium. Half of the sample was collected, while the remaining sample was treated with 0.6 mM H_2_O_2_ for 30 minutes at 30 °C at 200 rpm agitation. Cells were lysed by glass bead disruption agitation at 4°C in lysis buffer containing 50 mM Tris-acetate pH 7, 100 mM NaCl, 30 mM MgCl2, 20 mM iodoacetamide (IAM), and 1x complete mini EDTA-free protease inhibitor cocktail (Roche #04693159001). Cell extracts were clarified by centrifugation and were sedimented by ultracentrifugation for 120 min at 70,000 rpm (Beckman Optima Max-TL, TLA-110 rotor) at 4°C in a 50% sucrose cushion buffered in 50 mM Tris-acetate, pH 7.0, 150 mM NaCl, and 15 mM MgCl2. Ribosome pellets were resuspended in 250 μL digestion buffer (50 mM ACN pH 8, 50 mM NaCl, 8M Urea). Samples were diluted to < 1 M Urea with 50 mM ACN pH8, and protein concentration was determined by Bradford Assay (Bio-Rad, #5000205). 25 mg of resuspended ribosome pellet was digested with trypsin at a [20:1] protein: protease (w/w) ratio for 4 hours at 32°C. Digests were acidified to a final concentration of 1% FA for at least 15 minutes, clarified by centrifugation, and desalted by Sep-Pak C18 Cartridges (Waters #WAT051910). Samples were dried by lyophilization and resuspended in IAP buffer from PTMScan HS Ubiquitin/SUMO Remnant Motif (K-ε-GG) Kit (Cell Signaling #59322). K-GG containing heavy reference peptides were pooled and diluted into the IAP buffer resuspension to a final concentration of 1 fmol per peptide. Samples were incubated for 2 hours at 4°C with K-GG antibody beads and eluted with IAP Elution buffer (0.15% TFA) in accordance with the manufacturer’s protocol. Samples were again desalted by C18 Stage Tip cleanup prior to sample resuspension and targeted PRM or LC-MS/MS analysis.

#### Targeted PRM Mass Spectrometry analysis

The samples of K-GG-enriched peptides were reconstituted in 12 uL, and 3 uL of each was analyzed by LC-MS/MS using a Waters M-Class interfaced to a Thermo Exploris 480 MS/MS (n=5). Liquid chromatography utilized a trap-elute configuration and 90 min gradient. MS/MS used parallel reaction monitoring (targeted MS2) with a resolution of 30,000, AGC target of 200%, max IT of 200 ms and NCE of 30. The scheduled inclusion list used 4 min wide windows and a 3 s cycle time.

#### Label-free proteomics

The samples of K-GG-enriched peptides were analyzed by LC-MS/MS using a Waters M-Class interfaced to a Thermo Exploris 480 MS/MS (n=5). Liquid chromatography utilized a trap-elute configuration and 90 min gradient. MS/MS used data-independent acquisition (DIA) with a 120,000 resolution precursor scan from 375-1600 m/z, AGC target of 300% and max IT of 45 ms. DIA used 30,000 resolution, AGC target of 1000% and max IT of 60 ms a NCE of 30. DIA windows (20 total) spanned 400-1000 m/z with a window width of 30 m/z and window overlap of 1 m/z. Mass spectra were matched to previously built spectral library in Skyline and quantified.

#### Transition detection and peak integration with PRMKit

PRMkit is a new software package for processing parallel reaction monitoring (PRM) data in the context of targeted proteomics analysis (https://github.com/PRMkit). The core software performs feature detection on MS/MS data to extract ion chromatograms and determine the retention time positions of all potential fragment ions (transitions) for user-specified peptide sequences (target analytes). By learning patterns from all identified transitions within each peptide, the algorithm derives consensus start and end points of the ion chromatograms in each peptide and applies the rule during the peak integration of all its transitions. The user can easily visualize peak integration results using a custom script provided in the package. If desired, the user can manually change the integration boundaries of chosen analytes. PRMkit requires two input files: (a) a table containing the peptide sequences, precursor mass-to-charge ratios, and peptide charge states, and (b) a list of mzML files pointing to all analyses. The software produces a tab-delimited output table containing peak area values for individual transitions and pdf files for the ion chromatograms across all peptides and samples. For further downstream processing of the data, users can follow the example analysis demonstrated in the R markdown document in the Github repository.

### Dual luciferase reporter activity

Yeast strains transformed with Rluc-P2A-X-P2A-Fluc plasmids, where X represents a variable sequence, were grown to mid-log phase in SD complete medium. Cells were pelleted and resuspended in SD Methionine depleted (SD -Met) medium to induce plasmid expression for 60 minutes. Cells were then either left untreated or treated with 0.6 mM H_2_O_2_ or 4-NQO (1 μg/mL, 5 μg/mL) for an additional 30 minutes under agitation. Cells were then pelleted and disrupted by glass bead agitation at 4°C in 1x Passive Lysis Buffer provided in the Dual-Luciferase Reporter Assay System (Promega #E1910). Extracts were clarified by centrifugation, and protein concentration was determined by BCA assay (ThermoFisher #23225). Luminescence activities of Rluc and Fluc were collected for 3 ug of protein combined with respective substrates 10x diluted and were measured in a Spectra Max M3 (Molecular Devices) plate reader.

### Translation rate assays

The indicated SUB280-derived yeast strains were grown to mid-log phase in SD complete medium and back diluted to OD_600_ 0.1-0.2 in SD -Met medium. Cells were treated with 50 uM of HPG (L-Homopropargylglycine, Sigma, #900893) at OD_600_ 0.4-0.5. Cells were collected by centrifugation after 15, 30 and 45 min of HPG incubation at 30°C with agitation. For stress treatment, cells were incubated with 0.6 mM H_2_O_2_, 1 μg/mL 4-NQO, or 5 μg/mL 4-NQO for 15 min prior to HPG incubation for the times listed above. Pelleted cells were fixed overnight in 70% ethanol at 4°C. HPG conjugation with Alexa Fluor 488 was done with the Click-iT HPG Alexa Fluor Protein Synthesis Assay (ThermoFisher #C10428) per the manufacturer’s protocol. Alexa Fluor 488 fluorescent signal was measured in the BD FACS Symphony A1 flow cytometer using a 488 nm laser. FlowJo Software (Becton Dickinson) was used for single-cell population gates, histogram plots, and mean calculations.

### Gcn4-lacZ reporter assays

*GCN4-lacZ* expressing yeast cells from the indicated SUB280-derived strains were grown to mid-log phaser and disrupted by glass bead agitation at 4°C in buffer containing 1x PBS, 40 mM KCl, 10 mM MgCl_2_. Extracts were clarified by centrifugation and protein concentration was quantified by BCA assay (ThermoFisher #23225). 120 ug of protein was combined with substrate containing 15mM ONPG (2-Nitrophenyl-β-D-galactopyranoside, Goldbio, #N27510), 5mM DTT, 1x PBS, 40mM KCl, and 10mM MgCl2, and incubated for 60 min at 30°C. Absorbance was read at 420 nm in a Tecan Sunrise plate reader (Tecan # 30190079).

### Gcn4 target mRNA-seq analysis

RNA-seq data from WT and *hel2Δ* cells in the presence and absence of 0.1% MMS treatment (GSE150790) (Yan & Zaher, 2021) and from WT and *rad6Δ* cells in the presence and absence of 0.6 mM H_2_O_2_ treatment (GSE226082) (Meydan *et al*., 2023) were analyzed for differential expression of known targets of Gcn4 transcription regulation.

### Quantification and statistical analysis

Sample sizes and statistical tests used in this paper are described in the figure legends, and further details are provided in the methods section. All statistical analyses were performed on R (version 4.4.1) and RStudio (version 2024.09.0+375). Differences were considered statistically significant at a p-value < 0.05. For multiple comparison analyses, significant changes were calculated by Tukey’s test in conjunction with an ANOVA (p<0.05) or through multiple testing correction with Benjamini-Hochberg procedure.

---

**Figure S1.**
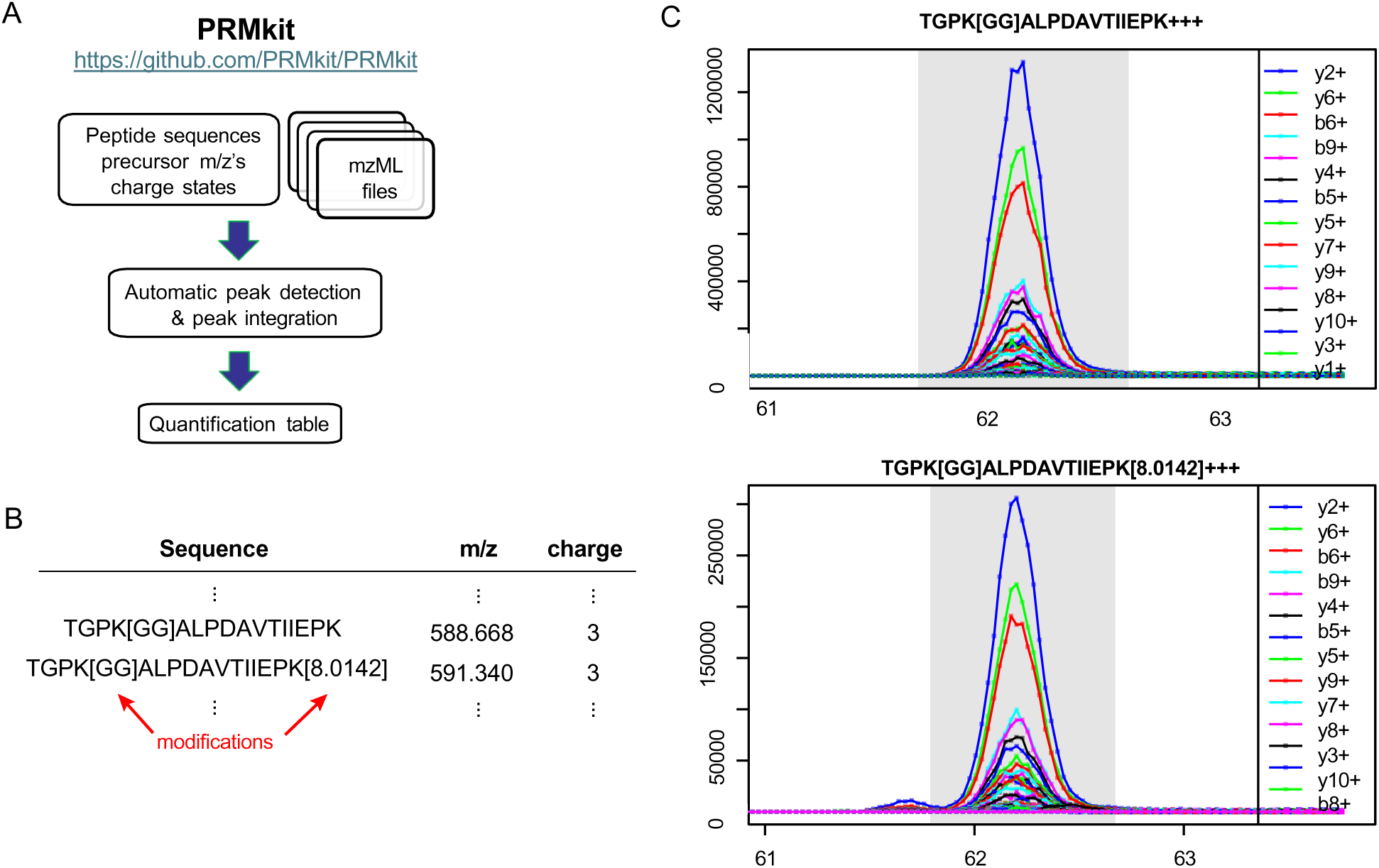
Quantification of peptide peak area integration by PRMkit. **(A)** Schematic of PRMkit chromatogram extraction and peak integration. **(B)** Example of peptide sequences, precursor m/z, and charge states required as input for PRMkit. **(C)** Example of PRMkit identification of the uS3.K200 biological (TGPK[GG]ALPDAVTIIEPK+++) and heavy-labeled (TGPK(GG)ALPDAVTIIEPK[8.0142]+++) peptide y and b ion transitions in decreasing order of integrated peak area. Gray rectangle shows the range of integration in each window.

**Figure S2.**
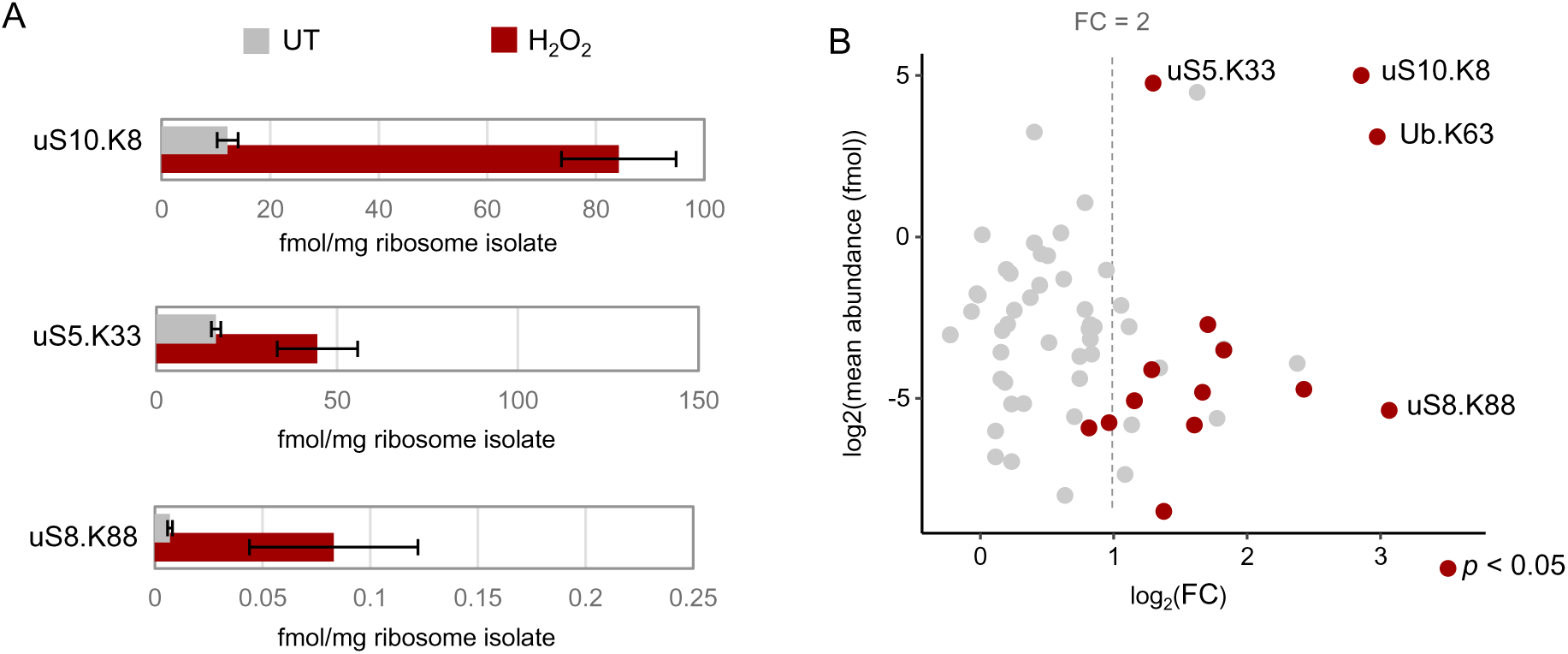
Quantification of key ribosome peptide abundance. **(A)** The abundance (fmol) of select ribosome site peptides per mg of ribosome from either untreated (UT) or 0.6 mM H_2_O_2_ treated (H_2_O_2_) is shown. Error bars represent SEM (n=5). **(B)** MA plot of the M(log ratio) and A(mean average) differences of Ub-modified ribosome sites in untreated and H_2_O_2_ conditions. Significance was calculated using a student’s paired *t* test, where * *p* < 0.05.

**Figure S3.**
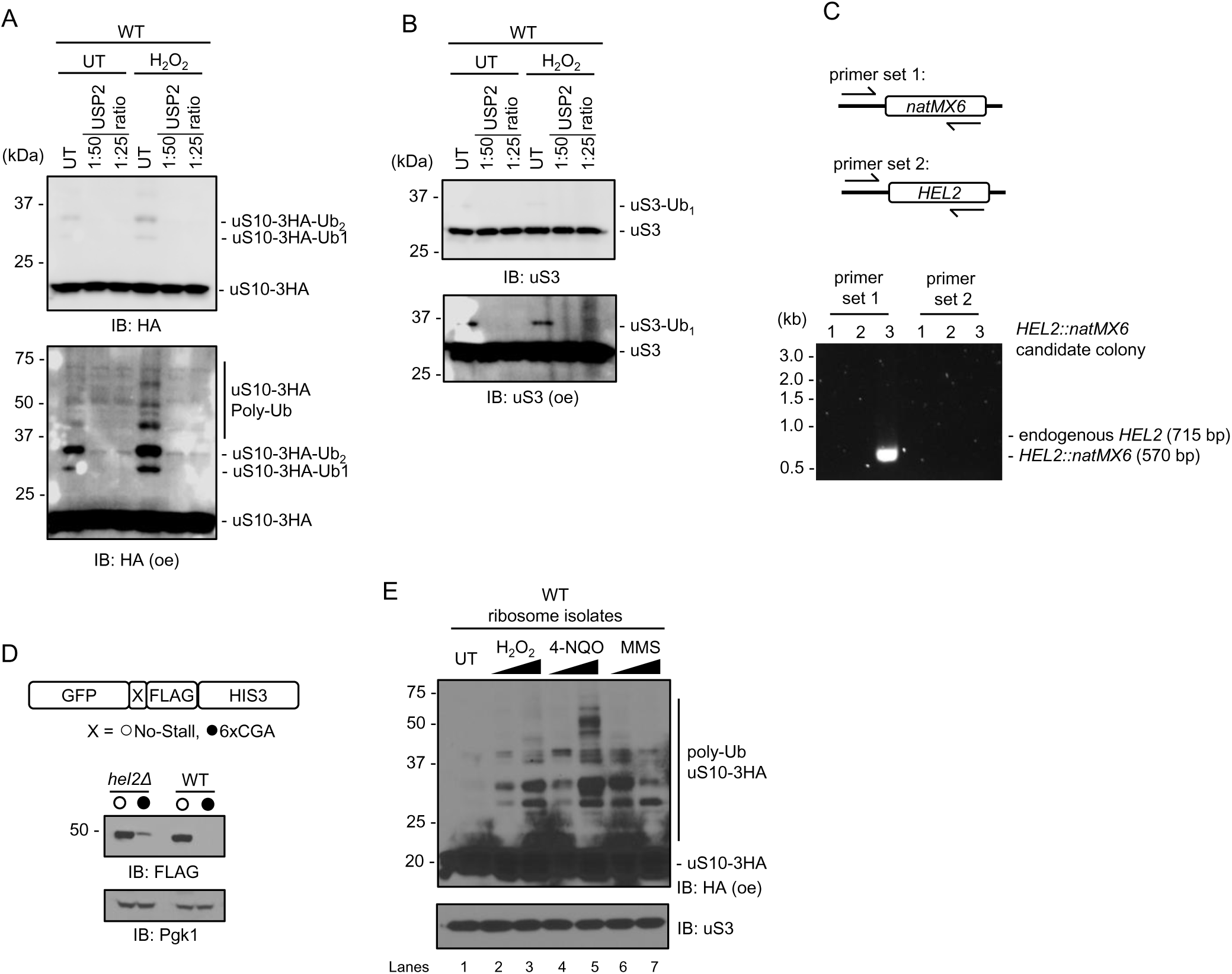
Dynamics of ribosomal protein ubiquitination in the absence of Hel2. **(A-B)** Western blot of polyubiquitin chain crash assay with USP2 deubiquitinase (1:50 or 1:25 ratio of DUB to protein). **(a)** anti-uS10-3HA, and **(b)** anti-uS3, from ribosome isolated from WT cells in untreated (UT) and 0.6 mM H_2_O_2_ treated conditions. **(C)** PCR confirmation of endogenous *HEL2* gene disruption with *natMX6*. Primer set 1 used to confirm insert of *natMX6* gene using forward primer aligning to upstream sequence of genomic *HEL2* and reverse primer that aligns internal to the *natMX6* sequence. Primer set 2 features forward primer aligning to upstream *HEL2* sequence and a reverse primer that aligns internal to the endogenous *HEL2* sequence. *Hel2::natMX6* candidate colony #3 confirmed to have disrupted *HEL2* gene. **(D)** Western blot for full-length GFP-X-FLAG-HIS3 reporter expression for the No-Stall and 6xCGA sequences in WT and *hel2Δ* with anti-FLAG. Anti-Pgk1 was used as a loading control. **(E)** Western blot for polyubiquitinated forms of uS10 in ribosome isolates from WT cells in untreated (UT), H_2_O_2_ (0.6 mM, 5 mM), 4-NQO (1 μg/mL, 5 μg/mL), and MMS (0.1%, 0.33%) treated conditions. Anti-uS3 used as loading control.

**Figure S4.**
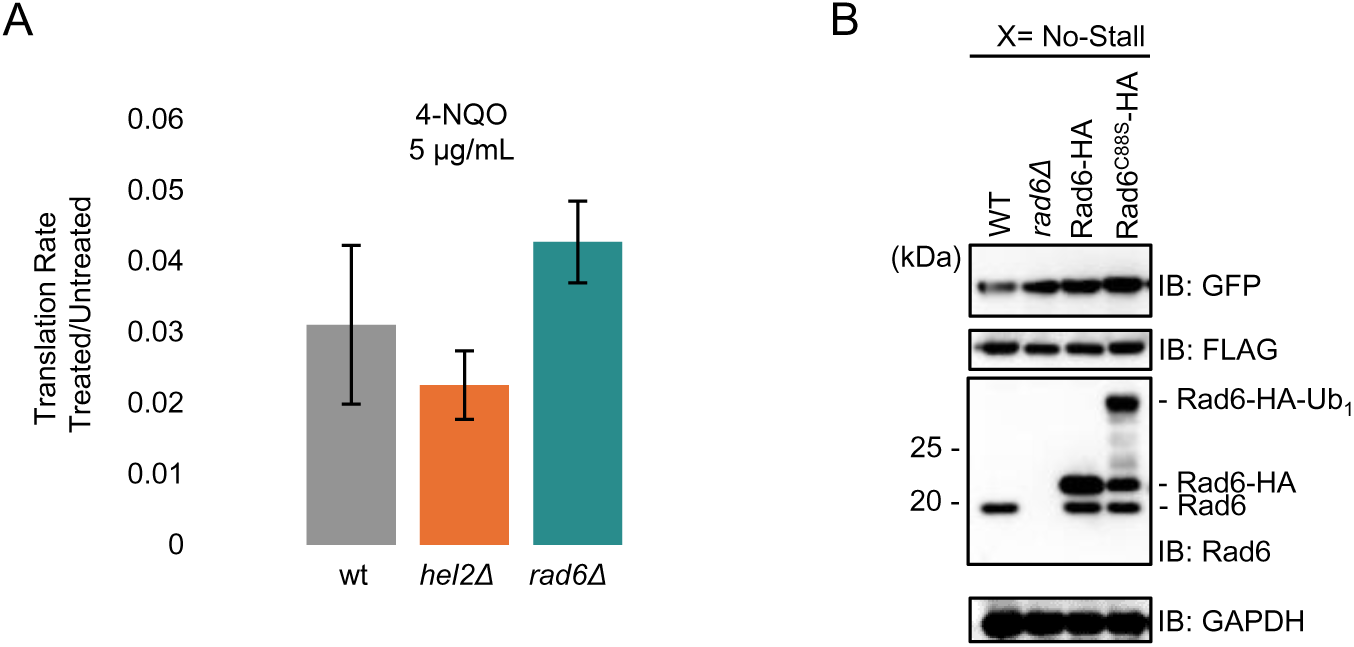
Translation rate determination and no-stalling reporter control. **(A)** Quantification of HPG incorporation upon exposure to 5 μg/mL 4-NQO in wild-type (WT), *hel2Δ*, and *rad6Δ*. Data were normalized to untreated levels. Error bars represent SEM (n=2). **(B)** Western blot for full-length GFP-No-Stall-FLAG-HIS3 reporter construct in WT, *rad6Δ,* and WT strains expressing HA-tagged Rad6 and Rad6 C88S mutant. Anti-GAPDH used as loading control.

## Notes

### Competing Interest Statement

The authors have declared no competing interest.

### Summary of Updates

All figures were revised and improved, new data were added and manuscript text was edited for clarity and content delivery

## REFERENCES

Advani VM, Ivanov P (2019) Translational Control under Stress: Reshaping the Translatome. BioEssays 41: 1900009

Aimiuwu J, Wang H, Chen P, Xie Z, Wang J, Liu S, Klisovic R, Mims A, Blum W, Marcucci G et al (2012) RNA-dependent inhibition of ribonucleotide reductase is a major pathway for 5-azacytidine activity in acute myeloid leukemia. Blood 119: 5229–5238

An H, Ordureau A, Körner M, Paulo JA, Harper JW (2020) Systematic quantitative analysis of ribosome inventory during nutrient stress. Nature 583: 303–309

Back S, Gorman AW, Vogel C, Silva GM (2019) Site-Specific K63 Ubiquitinomics Provides Insights into Translation Regulation under Stress. J Proteome Res 18: 309–318

Bailleul B, Daubersies P, Galiègue-Zouitina S, Loucheux-Lefebvre MH (1989) Molecular basis of 4-nitroquinoline 1-oxide carcinogenesis. Jpn J Cancer Res 80: 691–697

Barros GC, Guerrero S, Silva GM (2023) The central role of translation elongation in response to stress. Biochemical Society Transactions 51: 959–969

Bhuvanagiri M, Lewis J, Putzker K, Becker JP, Leicht S, Krijgsveld J, Batra R, Turnwald B, Jovanovic B, Hauer C et al (2014) 5&#x2010;azacytidine inhibits nonsense&#x2010;mediated decay in a MYC&#x2010;dependent fashion. EMBO Molecular Medicine 6: 1593–1609-1609

Blevins WR, Tavella T, Moro SG, Blasco-Moreno B, Closa-Mosquera A, Díez J, Carey LB, Albà MM (2019) Extensive post-transcriptional buffering of gene expression in the response to severe oxidative stress in baker’s yeast. Scientific Reports 9: 11005

Bowman GD, Poirier MG (2015) Post-translational modifications of histones that influence nucleosome dynamics. Chem Rev 115: 2274–2295

Buccitelli C, Selbach M (2020) mRNAs, proteins and the emerging principles of gene expression control. Nature Reviews Genetics 21: 630–644

Bustos D, Bakalarski CE, Yang Y, Peng J, Kirkpatrick DS (2012) Characterizing ubiquitination sites by peptide-based immunoaffinity enrichment. Mol Cell Proteomics 11: 1529–1540

Cavalieri V (2021) The Expanding Constellation of Histone Post-Translational Modifications in the Epigenetic Landscape. Genes (Basel) 12

Chen X, Shi C, He M, Xiong S, Xia X (2023) Endoplasmic reticulum stress: molecular mechanism and therapeutic targets. Signal Transduction and Targeted Therapy 8: 352

Costa-Mattioli M, Walter P (2020) The integrated stress response: From mechanism to disease. Science 368: eaat5314

Crawford RA, Pavitt GD (2019) Translational regulation in response to stress in. Yeast 36: 5–21

de Nadal E, Ammerer G, Posas F (2011) Controlling gene expression in response to stress. Nature Reviews Genetics 12: 833–845

Dever TE (1997) UsingGCN4as a Reporter of eIF2α Phosphorylation and Translational Regulation in Yeast. Methods 11: 403–417

Dougherty SE, Maduka AO, Inada T, Silva GM (2020) Expanding Role of Ubiquitin in Translational Control. Int J Mol Sci 21

Erpapazoglou Z, Walker O, Haguenauer-Tsapis R (2014) Versatile roles of k63-linked ubiquitin chains in trafficking. Cells 3: 1027–1088

Garshott DM, Sundaramoorthy E, Leonard M, Bennett EJ (2020) Distinct regulatory ribosomal ubiquitylation events are reversible and hierarchically organized. eLife 9: e54023

Gasch AP, Spellman PT, Kao CM, Carmel-Harel O, Eisen MB, Storz G, Botstein D, Brown PO (2000) Genomic Expression Programs in the Response of Yeast Cells to Environmental Changes. Molecular Biology of the Cell 11: 4241–4257

Gerashchenko MV, Lobanov AV, Gladyshev VN (2012) Genome-wide ribosome profiling reveals complex translational regulation in response to oxidative stress. Proceedings of the National Academy of Sciences 109: 17394–17399

Ghulam MM, Catala M, Abou Elela S (2020) Differential expression of duplicated ribosomal protein genes modifies ribosome composition in response to stress. Nucleic Acids Research 48: 1954–1968

Hinnebusch AG (2005) TRANSLATIONAL REGULATION OF GCN4 AND THE GENERAL AMINO ACID CONTROL OF YEAST*. Annual Review of Microbiology 59: 407–450

Holcik M, Sonenberg N (2005) Translational control in stress and apoptosis. Nature Reviews Molecular Cell Biology 6: 318–327

Inada T (2020) Quality controls induced by aberrant translation. Nucleic Acids Research 48: 1084–1096

Iyer KV, Müller M, Tittel LS, Winz M-L (2023) Molecular Highway Patrol for Ribosome Collisions. ChemBioChem 24: e202300264

Jomova K, Raptova R, Alomar SY, Alwasel SH, Nepovimova E, Kuca K, Valko M (2023) Reactive oxygen species, toxicity, oxidative stress, and antioxidants: chronic diseases and aging. Archives of Toxicology 97: 2499–2574

Kettenbach AN, Rush J, Gerber SA (2011) Absolute quantification of protein and post-translational modification abundance with stable isotope–labeled synthetic peptides. Nature Protocols 6: 175–186

Khoddami V, Cairns BR (2013) Identification of direct targets and modified bases of RNA cytosine methyltransferases. Nature Biotechnology 31: 458–464

Kirkpatrick DS, Denison C, Gygi SP (2005) Weighing in on ubiquitin: the expanding role of mass-spectrometry-based proteomics. Nature Cell Biology 7: 750–757

Kito K, Ito T (2008) Mass spectrometry-based approaches toward absolute quantitative proteomics. Curr Genomics 9: 263–274

Knight JRP, Garland G, Pöyry T, Mead E, Vlahov N, Sfakianos A, Grosso S, De-Lima-Hedayioglu F, Mallucci GR, von der Haar T et al (2020) Control of translation elongation in health and disease. Disease Models & Mechanisms 13

Kohda K, Tada M, Kasai H, Nishimura S, Kawazoe Y (1986) Formation of 8-hydroxyguanine residues in cellular DNA exposed to the carcinogen 4-nitroquinoline 1-oxide. Biochem Biophys Res Commun 139: 626–632

Kosower NS, Kosower EM (1995) [11] Diamide: An oxidant probe for thiols. In: *Methods in Enzymology*, pp. 123-133. Academic Press:

Li LH, Olin EJ, Buskirk HH, Reineke LM (1970) Cytotoxicity and mode of action of 5-azacytidine on L1210 leukemia. Cancer Res 30: 2760–2769

Liu P, Gan W, Su S, Hauenstein AV, Fu T-m, Brasher B, Schwerdtfeger C, Liang AC, Xu M, Wei W (2018) K63-linked polyubiquitin chains bind to DNA to facilitate DNA damage repair. Science Signaling 11: eaar8133

Llácer Jose L, Hussain T, Marler L, Aitken Colin E, Thakur A, Lorsch Jon R, Hinnebusch Alan G, Ramakrishnan V (2015) Conformational Differences between Open and Closed States of the Eukaryotic Translation Initiation Complex. Molecular Cell 59: 399–412

Lushchak VI (2014) Free radicals, reactive oxygen species, oxidative stress and its classification. Chemico-Biological Interactions 224: 164–175

MacLean B, Tomazela DM, Shulman N, Chambers M, Finney GL, Frewen B, Kern R, Tabb DL, Liebler DC, MacCoss MJ (2010) Skyline: an open source document editor for creating and analyzing targeted proteomics experiments. Bioinformatics 26: 966–968

Madiraju C, Novack JP, Reed JC, Matsuzawa SI (2022) K63 ubiquitination in immune signaling. Trends Immunol 43: 148–162

Manohar S, Jacob S, Wang J, Wiechecki KA, Koh HWL, Simões V, Choi H, Vogel C, Silva GM (2019) Polyubiquitin Chains Linked by Lysine Residue 48 (K48) Selectively Target Oxidized Proteins In Vivo. Antioxidants & Redox Signaling 31: 1133–1149

Martín-Villanueva S, Gutiérrez G, Kressler D, de la Cruz J (2021) Ubiquitin and ubiquitin-like proteins and domains in ribosome production and function: chance or necessity? International Journal of Molecular Sciences 22: 4359

Martínez-Férriz A, Ferrando A, Fathinajafabadi A, Farràs R (2022) Ubiquitin-mediated mechanisms of translational control. Seminars in Cell & Developmental Biology 132: 146–154

Martins D, English AM (2014) Catalase activity is stimulated by H(2)O(2) in rich culture medium and is required for H(2)O(2) resistance and adaptation in yeast. Redox Biol 2: 308–313

Mascarenhas C, Edwards-Ingram LC, Zeef L, Shenton D, Ashe MP, Grant CM (2008) Gcn4 Is Required for the Response to Peroxide Stress in the Yeast Saccharomyces cerevisiae. Molecular Biology of the Cell 19: 2995–3007

Matsuki Y, Matsuo Y, Nakano Y, Iwasaki S, Yoko H, Udagawa T, Li S, Saeki Y, Yoshihisa T, Tanaka K (2020) Ribosomal protein S7 ubiquitination during ER stress in yeast is associated with selective mRNA translation and stress outcome. Scientific reports 10: 19669

Matsuo Y, Ikeuchi K, Saeki Y, Iwasaki S, Schmidt C, Udagawa T, Sato F, Tsuchiya H, Becker T, Tanaka K et al (2017) Ubiquitination of stalled ribosome triggers ribosome-associated quality control. Nature Communications 8: 159

Matsuo Y, Inada T (2023) Co-Translational Quality Control Induced by Translational Arrest. Biomolecules 13: 317

Matsuo Y, Uchihashi T, Inada T (2023) Decoding of the ubiquitin code for clearance of colliding ribosomes by the RQT complex. Nature Communications 14: 79

Meng EC, Goddard TD, Pettersen EF, Couch GS, Pearson ZJ, Morris JH, Ferrin TE (2023) UCSF ChimeraX: Tools for structure building and analysis. Protein Science 32: e4792

Meydan S, Barros GC, Simões V, Harley L, Cizubu BK, Guydosh NR, Silva GM (2023) The ubiquitin conjugase Rad6 mediates ribosome pausing during oxidative stress. Cell Reports 42

Monem PC, Arribere JA (2024) A ubiquitin language communicates ribosomal distress. Seminars in Cell & Developmental Biology 154: 131–137

Morano KA, Grant CM, Moye-Rowley WS (2012) The Response to Heat Shock and Oxidative Stress in Saccharomyces cerevisiae. Genetics 190: 1157–1195

Nanjaraj Urs AN, Lasehinde V, Kim L, McDonald E, Yan LL, Zaher HS (2024) Inability to rescue stalled ribosomes results in overactivation of the integrated stress response. Journal of Biological Chemistry 300

Ordureau A, Paulo JA, Zhang W, Ahfeldt T, Zhang J, Cohn EF, Hou Z, Heo JM, Rubin LL, Sidhu SS et al (2018) Dynamics of PARKIN-Dependent Mitochondrial Ubiquitylation in Induced Neurons and Model Systems Revealed by Digital Snapshot Proteomics. Mol Cell 70: 211–227.e218

Pakos-Zebrucka K, Koryga I, Mnich K, Ljujic M, Samali A, Gorman AM (2016) The integrated stress response. EMBO Rep 17: 1374–1395

Peng J, Schwartz D, Elias JE, Thoreen CC, Cheng D, Marsischky G, Roelofs J, Finley D, Gygi SP (2003) A proteomics approach to understanding protein ubiquitination. Nature Biotechnology 21: 921–926

Peterson AC, Russell JD, Bailey DJ, Westphall MS, Coon JJ (2012) Parallel reaction monitoring for high resolution and high mass accuracy quantitative, targeted proteomics. Mol Cell Proteomics 11: 1475–1488

Postnikoff SDL, Johnson JE, Tyler JK (2017) The integrated stress response in budding yeast lifespan extension. Microb Cell 4: 368–375

Rahman S, Wolberger C (2024) Breaking the K48-chain: linking ubiquitin beyond protein degradation. Nature Structural & Molecular Biology 31: 216–218

Santos CM, Cizubu BK, Okonkwo DA, Chen C-Y, Maske N, Snyder NA, Simões V, Washington EJ, Silva GM (2024) Redox control of the deubiquitinating enzyme Ubp2 regulates translation during stress. Journal of Biological Chemistry 300: 107870

Schneider CA, Rasband WS, Eliceiri KW (2012) NIH Image to ImageJ: 25 years of image analysis. Nature Methods 9: 671–675

Sheng X, Xia Z, Yang H, Hu R (2024) The ubiquitin codes in cellular stress responses. Protein & Cell 15: 157–190

Shenton D, Smirnova JB, Selley JN, Carroll K, Hubbard SJ, Pavitt GD, Ashe MP, Grant CM (2006) Global Translational Responses to Oxidative Stress Impact upon Multiple Levels of Protein Synthesis*. Journal of Biological Chemistry 281: 29011–29021

Silva GM, Finley D, Vogel C (2015) K63 polyubiquitination is a new modulator of the oxidative stress response. Nature Structural & Molecular Biology 22: 116–123

Simms CL, Yan LL, Zaher HS (2017) Ribosome Collision Is Critical for Quality Control during No-Go Decay. Molecular Cell 68: 361–373.e365

Simms CL, Zaher HS (2016) Quality control of chemically damaged RNA. Cellular and Molecular Life Sciences 73: 3639–3653

Simões V, Cizubu BK, Harley L, Zhou Y, Pajak J, Snyder NA, Bouvette J, Borgnia MJ, Arya G, Bartesaghi A et al (2022) Redox-sensitive E2 Rad6 controls cellular response to oxidative stress via K63-linked ubiquitination of ribosomes. Cell Reports 39

Taymaz-Nikerel H, Cankorur-Cetinkaya A, Kirdar B (2016) Genome-Wide Transcriptional Response of Saccharomyces cerevisiae to Stress-Induced Perturbations. Front Bioeng Biotechnol 4: 17

Thannickal VJ, Fanburg BL (2000) Reactive oxygen species in cell signaling. American Journal of Physiology-Lung Cellular and Molecular Physiology 279: L1005–L1028

van Bentum M, Selbach M (2021) An Introduction to Advanced Targeted Acquisition Methods. Molecular & Cellular Proteomics 20

Vind AC, Genzor AV, Bekker-Jensen S (2020) Ribosomal stress-surveillance: three pathways is a magic number. Nucleic Acids Res 48: 10648–10661

Vogel C, Silva GM, Marcotte EM (2011) Protein Expression Regulation under Oxidative Stress*. Molecular & Cellular Proteomics 10: M111.009217

Williams TD, Rousseau A (2024) Translation regulation in response to stress. The FEBS Journal n/a

Wilson DN, Doudna Cate JH (2012) The structure and function of the eukaryotic ribosome. Cold Spring Harb Perspect Biol 4

Winterbourn CC (2008) Reconciling the chemistry and biology of reactive oxygen species. Nature Chemical Biology 4: 278–286

Wong C-M, Ching Y-P, Zhou Y, Kung H-F, Jin D-Y (2003) Transcriptional regulation of yeast peroxiredoxin gene TSA2 through Hap1p, Rox1p, and Hap2/3/5p. Free Radical Biology and Medicine 34: 585–597

Wong C-M, Zhou Y, Ng RWM, Kung H-f, Jin D-Y (2002) Cooperation of Yeast Peroxiredoxins Tsa1p and Tsa2p in the Cellular Defense against Oxidative and Nitrosative Stress *. Journal of Biological Chemistry 277: 5385–5394

Wu CC-C, Peterson A, Zinshteyn B, Regot S, Green R (2020) Ribosome Collisions Trigger General Stress Responses to Regulate Cell Fate. Cell 182: 404–416.e414

Xu P, Duong DM, Seyfried NT, Cheng D, Xie Y, Robert J, Rush J, Hochstrasser M, Finley D, Peng J (2009) Quantitative Proteomics Reveals the Function of Unconventional Ubiquitin Chains in Proteasomal Degradation. Cell 137: 133–145

Yan LL, Simms CL, McLoughlin F, Vierstra RD, Zaher HS (2019) Oxidation and alkylation stresses activate ribosome-quality control. Nature Communications 10: 5611

Yan LL, Zaher HS (2021) Ribosome quality control antagonizes the activation of the integrated stress response on colliding ribosomes. Molecular Cell 81: 614–628.e614

Zhou Y, Kastritis PL, Dougherty SE, Bouvette J, Hsu AL, Burbaum L, Mosalaganti S, Pfeffer S, Hagen WJH, Förster F et al (2020) Structural impact of K63 ubiquitin on yeast translocating ribosomes under oxidative stress. Proceedings of the National Academy of Sciences 117: 22157–22166

